# Ultra-high field microstructural MRI of living cortical organoids

**DOI:** 10.64898/2026.04.19.719203

**Authors:** Tatiana Nikolaeva, Channa E. Jakobs, Maxime Yon, Youri Adolfs, Rico Singer, R. Jeroen Pasterkamp, Julia R. Krug, Chantal M.W. Tax

**Affiliations:** Center for Image Sciences, University Medical Center Utrecht (UMCU), Utrecht, The Netherlands; Laboratory of BioNanoTechnology, Wageningen University & Research (WUR), Wageningen, The Netherlands; Department of Translational Neuroscience, University Medical Center Brain Center, Utrecht University (UU), Utrecht, The Netherlands; Laboratoire Traitement du Signal et de l’Image (LTSI), Université de Rennes, Rennes, France; ICube Laboratory, University of Strasbourg, CNRS, Strasbourg, France; Cardiff University Brain Research Imaging Centre (CUBRIC), School of Physics and Astronomy, Cardiff University, Cardiff, United Kingdom

**Author notes:** authors with equal contributions (Julia R. Krug, Chantal M.W. Tax).

## Abstract

Quantitative microstructural magnetic resonance imaging (MRI) can noninvasively characterize tissue configuration at micrometer scales, but clinical uptake is limited by validation and optimization in human-relevant scenarios. Organoids are powerful human-relevant tissue models, yet translation is hampered by lack of non-destructive, longitudinal microstructural assessment. Bridging these gaps, microstructural MRI of living organoids can accelerate MRI biomarker and organoid development and validation. Here, we address key obstacles to enable organoid microstructural MRI. First, we use a unique 28.2 T MRI system to achieve spatial resolution with adequate signal-to-noise ratio and feasible scan times. Second, we implement flexible acquisitions with fast readouts to expand multivariate experimental capacity. Third, we develop a workflow combining 3D MRI and 3D lightsheet microscopy for cross-modality anatomical comparison beyond 2D. Using this platform, we demonstrate microstructural MRI of cortical organoids with resolutions down to (20 *µm*)^3^, revealing anisotropy, heterogeneity, maturation-dependent differences, and temporal changes in cortical organoids. Correlative lightsheet microscopy confirms correspondence to axonal and nuclear architecture. This platform enables live-organoid MRI as a complementary tool to human- and animal imaging for robust microstructural assessment.

## 1 Introduction

Non-invasive and non-destructive imaging techniques of biological tissues are essential to study structure and function without disrupting their 3D architecture and metabolic processes. In this context, magnetic resonance imaging (MRI) can provide detailed 3D characterization over time without ionizing radiation, making it uniquely suited for clinical and research applications in both whole organisms or organs [1] and tissue- or cell cultures. Importantly, as opposed to invasive and/or destructive imaging, MRI can facilitate the translation of findings between *in vivo, ex vivo*, and *in vitro* systems. Diffusion MRI (dMRI) is a specialized MRI technique with which the MRI signal is modulated by tissue microstructure on the cellular scale [2, 3]. Consequently, dMRI can provide characterization of structural and histological tissue properties, including fraction, size, shape, and orientation of spaces within and around cells. It does so by applying additional magnetic field gradients that sensitize the measurement to the diffusion of water molecules, which probe their spatial environment in the order of micrometers during the typical time scale of MRI experiments [4]. Relaxation MRI (*T*_1_- and *T*_2_-relaxation) can provide additional and complementary microstructural information on compartment sizes [5, 6], and myelination [7, 8], among others. Microstructural MRI is a quantitative technique that combines multiple diffusion- and/or relaxation-weighted measurements – obtained by varying the MRI-sequence timings and gradient-waveform shapes, directions, and strengths – with mathematical modelling to extract salient signal- or biophysical features related to tissue microstructure [9]. It has been used extensively to quantitatively study tissue microstructure in both healthy states and numerous diseases. Organoids are rapidly emerging as powerful, 3D *in vitro* models that aim to recapitulate key structural and functional features of human tissues, with broad implications for developmental biology, disease modeling and replacement of animal experiments [10]. Microstructural MRI of organoids has the potential to push the boundaries of both current MRI methodology and organoid research, as well as offer a powerful translational pipeline between clinical imaging and fundamental biology. Specifically, for MRI research, it could provide a complementary platform to current human and animal MRI research. Furthermore, microstructural MRI of organoids can address key issues in understanding, validating, and developing MRI markers. Specifically, by mitigating structural changes and mislocalizations caused by invasive removal of tissue from living organisms, improved structural correspondence could be achieved between MRI and subsequent microscopy in organoids. For organoid research, MRI could provide a complementary window into investigating organoid structure in addition to optical microscopy [11] and electron microscopy [12] without the need for fixing, clearing, staining, or slicing. Finally, comparing MRI markers in organoids and humans could lead to the development of more physiologically relevant *in vitro* models for human tissue, ultimately accelerating the development of diagnostics and therapies. MRI studies have shown the feasibility of imaging small *in vitro* models such as spheroids [13–20] and organoids ([21–25]). Notably, MRI-contrast agents were used in some cases on fixed organoids to shorten the experiment time for high spatial resolutions [21, 22]. Qualitative MRI based on relaxation-weighting at an MRI field strength of 3 Tesla (*T*) has been used to determine the volume of embedded cortical organoids in sodium alginate and gelatin [23] (resolution (300 *µm*)^3^, measurement time 7 *h* 30 *min*) and to visualize amyloid plaques in fixed brain cortical organoids (11.7 *T* with cryoprobe, resolution (15 *µm*)^3^, measurement time 28 *h* 30 *min*) [22]. Recent work has demonstrated the feasibility of dMRI on organoids at 11.7 *T*, either fixed in paraformaldehyde with high resolution ((30 *µm*)^3^, measurement time 35 *h* [21], 1500 *mT/m* gradients) or kept in agarose at low resolution (310 × 310 × 800 *µm*^3^, 1 *h* 25 *min*, 1500 *mT/m* gradients [25]). dMRI together with an AI-based image analysis pipeline has been shown to be promising for monitoring the development of organoids (9.4 *T*, (80 *µm*)^3^, 15 *min* 40 *s* and 100 × 100 × 1500 *µm*^3^, 23 *min* 5 *s*) [24]. Advanced dMRI approaches using so-called oscillating gradient waveforms were able to distinguish different cell sizes in spheroids embedded in agarose gel-solidified culture medium (9.4 *T*, 80 × 80 × 600 *µm*^3^, measurement time 1 *h* 7 *min*, 700 *mT/m* gradients) [15].

While the promise of using MRI for organoid imaging has been shown in a handful of recent studies, key challenges limit the ability to unlock the full potential of microstructural MRI in organoids and its comparison to subsequent microscopy. The large typical voxel sizes in (pre-)clinical MRI systems that have been used so far (100-1000 *µm*) make it challenging to resolve spatial heterogeneity within small organoids and cause a large spatial scale gap with the area covered by optical- and electron microscopy, in turn hampering accurate spatial matching and thus validation of MRI markers. An increased spatial resolution can be achieved by acquiring and averaging multiple MRI measurements to increase signal-to-noise ratio (SNR), but this drastically increases scan time, which makes imaging of living organoids challenging. This is further exacerbated by the need to acquire multiple MRI images with different acquisition settings for microstructural MRI. Fixation or embedding organoids can extend the time window for MRI measurements, whereas contrast agents can shorten measurement time for a single image, but these procedures could cause functional and structural changes compared to *in vivo* (e.g. cell size, intrinsic diffusion, and membrane integrity and permeability) [26], and/or cause changes in MRI-observed diffusion markers [27, 28].

The current work aims to address key challenges for extensive microstructural MRI of living brain organoids [29] by increasing the SNR while maintaining acceptable acquisition times for *in vivo* imaging. This is achieved by using the state-of-the-art highest field strength for MRI (28.2 *T*) and small-diameter (5 *mm*) radiofrequency (RF) coils to obtain a high SNR per unit time without the use of contrast agent. The 28.2 *T* system furthermore includes a triaxial gradient insert which enables gradient field strengths up to 3 *T/m* per axis with a slew rate of 30,000 *T/m/s* for efficient diffusion-encoding. Combined with the implementation and evaluation of fast echo planar image (EPI) readout strategies, this provides an optimal measurement setup for detailed and efficient microstructural MRI of living non-embedded and non-fixed organoids in liquid medium. The setup is used [30, 31] to investigate cortical organoid microstructure with a variety of diffusion encodings to study changes over extended measurement times, demonstrate structural heterogeneity and anisotropy across the organoid, and showcase the feasibility of detecting differences in diffusion-restriction between young and mature organoids. Finally, this work develops and implements a workflow for subsequent 3D MRI and 3D lightsheet (LS) microscopy of intact cortical organoids. Moving beyond qualitative comparison of 3D MRI with 2D microscopy slices in previous work [21–23, 25, 32], the proposed workflow can ultimately be used for 3D quantitative comparison of histological markers derived from both modalities.

## 2 Results and Discussion

This section will first present results on the implementation of the 3D MRI and microscopy workflow (Section 2.1), and will subsequently describe the evaluation of fast and high-resolution EPI image readout techniques at 28.2 *T* (Section 2.2). Using this setup, Section 2.3 will describe the results of a range of microstructural MRI experiments performed on cortical organoids and discuss the findings in light of previous literature.

### 2.1 Workflow for microstructural MRI of living organoids with subsequent lightsheet microscopy

The developed workflow for living brain organoids is depicted schematically in Figure 1 (a). The human-induced pluripotent stem cell (hiPSC) derived cortical brain organoids were transferred to MRI tubes filled with culturing medium for subsequent imaging at 37 °*C* on a 28.2 *T* vertical bore MRI scanner. Spin-echo (SE) MRI protocols were implemented with a variety of timings to modulate relaxation-contrast (echo time *TE* and repetition time *TR*, or respective delays *τ*_*E*+_ and *τ*_*R*+_) and diffusion encodings to modulate diffusion-contrast (trapezoidal waveforms with gradient-lobe duration *δ*, gradient-lobe separation Δ, gradient strength **g** and *b*-value; and arbitrary gradient waveforms, Figure 1 (b), Table (1)). Quantitative MRI signal modelling and assessment yielded information on organoid microstructure (see Section 2.3).

**Figure 1:**
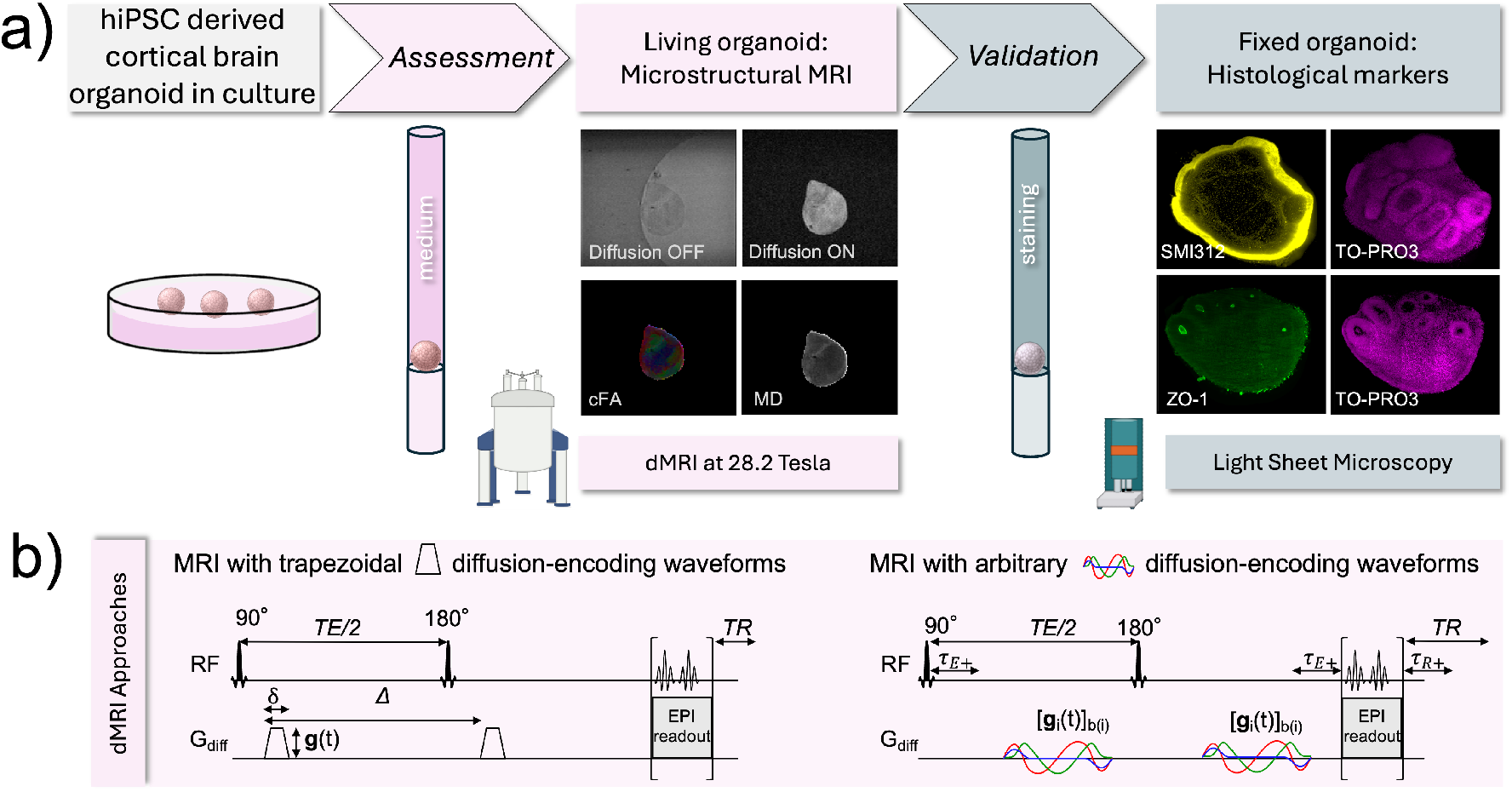
(a) Schematic of the developed workflow for microstructural imaging of living cortical brain organoids. An organoid in medium (non-fixed and non-embedded) undergoes microstructural 3D MRI assessment, including diffusion- (dMRI) and relaxation-weighted MRI. Subsequently, the organoid is fixed in the MR tube and immunostained for SMI312 (axons), TO-PRO3 (nuclei), and ZO-1 (ventricular-like zones (VLZs)). Subsequently, 3D lightsheet (LS) microscopy assessment is performed. Shown are examples of dMRI images: without and with diffusion weighting, derived color-coded macroscopic fractional anisotropy (cFA) and mean diffusivity (MD); and LS images stained for: 1) axons/ nuclei (top row) and 2) VLZs/nuclei (bottom row). (b) Two MRI sequences with fast Echo-Planar Imaging (EPI) image readout, and trapezoidal (left) and arbitrary (right) diffusion-encoding gradient waveforms, are schematically represented. The trapezoidal waveforms are presented with duration *δ*, strength *g*, a gradient-lobe separation Δ, echo time (*TE*) and repetition time (*TR*). The arbitrary waveforms 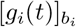 have variable shapes *g*_*i*_(*t*) and diffusion weighting (*b*-value *b*_*i*_ [*ms/µm*^2^]), and each measurement can have variable echo time delays (*τ*_*E*+_) and repetition time delays (*τ*_*R*+_) to achieve a given *TE* and *TR* without changing the diffusion-encoding timings.

**Table 1:** dMRI experiments and their main image acquisition parameters for microstructural assessment of cortical brain organoids. The young and mature organoids evaluated in this work are listed at the bottom of the table, alongside the MRI experiments in which they were used. The color coding is consistent with the figures in this work. 1 and 2 refer to immunostaining with ZO-1 and TO-PRO3 (1), or SMI312 and TO-PRO3 (2). * indicates that for Young 08 organoid, the protocol for the diffusion tensor images was slightly adjusted: 3D PGSE EPI, segments 3, TE/TR 10.5/3500 *ms*, resolution (30 *µm*)^3^, FOV (4.2 *mm*)^3^.

| Experiment | Longitudinal ADC changes |  | Diffusion kurtosis | ADC diffusion-time dependence | Diffusion tensor imaging/ Tractography | D(w)-R<sub>1</sub>-R<sub>2</sub> |
| --- | --- | --- | --- | --- | --- | --- |
| MR Method | 2D PGSE EPI |  | 3D PGSE EPI | 3D PGSE EPI | 3D PGSE EPI | 3D DOR_R1_R2_EPI |
| Segments | 8 |  | 8 | 8 | 8 | 1 |
| b-value [ms/μm<sup>2</sup>] | 0.0 / 1.0 |  | 0.0 / 0.6 / 1.2 / 1.8 / 2.4 / 3.0 | 0.0 / 0.25 / 0.4 / 1.0 | 0 / 1.0 | 0.3 – 4.07 |
| directions | 3 |  | 1 | 1 | 9 | 1 – 12 |
| δ [ms] | 1.0 |  | 1.2 | 0.9 | 0.9 | 3 – 9 |
| Δ [ms] | 3.0 |  | 4.3 | 2.73 / 5 / 10 / 15 / 25 | 2.73 | Not relevant |
| TE [ms] | 8.1 | 13 | 9 | 29.5 | 9.4 | 16 – 46 |
| TR [ms] | 3800 | 3800 | 3800 | 3800 | 3800 | 920 – 3430 |
| Resolution [μm<sup>3</sup>] | 40.6x40.6x120 | 44.8x44.8x120 | (44.8)<sup>3</sup> | (44.8)<sup>3</sup> | (19.5)<sup>3</sup> | (50)<sup>3</sup> |
| FOV [mm<sup>3</sup>] | 2.6 x 2.6 | 4.3 x 4.3 | (4.3)<sup>3</sup> | (4.3)<sup>3</sup> | (2.5)<sup>3</sup> | (2.55)<sup>3</sup> |
| Tube size [mm] | 3 | 5 | 5 | 5 | 3 | 3 |
| Measurement time | 8 min | 8 min | 4 h 51 min | 12 h 58 min | 8 h 10 min | 15 h 18 min |
| Organoids (age) | Young 02(D69)<sup>2</sup> | Young 03(D74)<sup>1</sup><br>Mature 01(D253)<sup>2</sup> | Young 03(D74)<sup>1</sup><br>Young 04(D76)<sup>1</sup><br>Young 05(D75)<sup>1</sup><br>Mature 01(D253)<sup>2</sup> | Young 06(D70)<sup>2</sup><br>Mature 02(D255)<sup>1</sup> | Young 01(D72)<sup>2</sup><br>Young 07(D70)<sup>1</sup><br>Young 08(D73)*<sup>2</sup> | Young 09(D72)<sup>1</sup> |

After MRI acquisition, organoids were fixed for further immunostaining with TO-PRO3 to visualize nuclei (Table (1)). Additionally, some organoids were immunostained with ZO-1 to identify ventricular-like zones (VLZs) and with SMI312 to visualize axons (Table (1)). All immunostained organoids were analyzed using LS microscopy for 3D visualization and correlation with MRI. In this work, a total of 25 cortical brain organoids were examined. 12 organoids were designated for dMRI sequence optimization, two of which were examined while fixed. Ultimately, 13 organoids completed the developed pipeline, which included a microstructural MRI experiment followed by immunostaining and LS microscopy imaging. 17 organoids out of 25 underwent the immunostaining procedure, while 4 were lost or significantly damaged during the staining process. Among the 13 cortical organoids studied, 10 organoids were young (ranging in age from day D69 to D76), and 3 were more mature (ranging from D253 to D255).

This workflow allows for comparing microstructural MRI markers on living brain organoids with histological markers from LS microscopy on the same organoid specimen after fixation, by manual alignment on structures like VLZs and spatial patterns in dMRI and microscopy maps. Figure 2 shows dMRI images (with a single direction of diffusion encoding) and LS images of young and mature cortical organoids, clearly revealing microstructural variability across the organoids. A qualitative comparison suggests that dMRI images reflect areas of diffusion restriction and anisotropy. For example, around VLZs in the young organoid dMRI shows lower signal intensity where anisotropic structures radially point outward in the direction of the encoding gradient and higher intensity where they are perpendicular. Corresponded LS images reveal high nuclear density in these areas. Nuclear density estimates of organoids on Figure 2 – obtained from nuclear counts on the nuclei-stained 3D LS images and organoid volume estimates on 3D dMRI images – suggested overall higher nuclear density in young organoid (0.00068 *nuclei/µm*^3^) compared to the mature one (0.00045 *nuclei/µm*^3^). This trend was consistent across other organoids (not shown).

**Figure 2:**
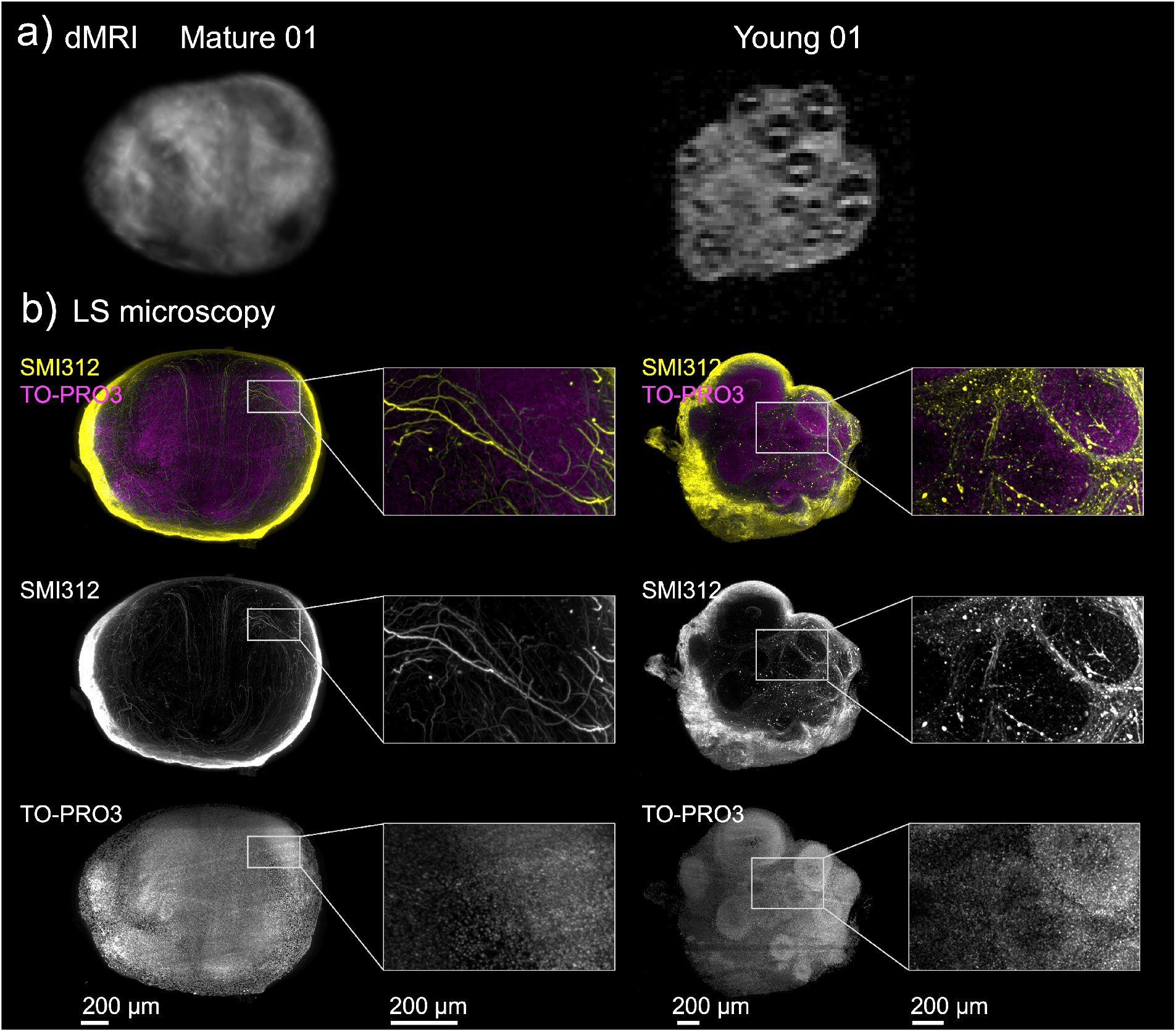
(a) Diffusion-weighted images (b-value 1 *ms/µm*^2^) of mature 01 and young 01 cortical brain organoids. (b) 2D visualization of immunostained organoids with antibodies recognizing SMI312 (axons) and TO-PRO3 (nuclei). 3D image obtained by subsequent lightsheet (LS) microscopy. For visualization purposes, the slice thickness of the LS images was 200 *µm*.

The choice of staining options in the current study was constrained by the four–channel LS microscopy setup. Future work will incorporate a broader range of stainings (e.g. different cell types and membranes) and investigate (additional) confocal and electron microscopy for further microstructural correlation. A remaining challenge is the co-registration of the MRI and LS microscopy images for further quantitative spatial matching and validation of MRI features. Non-rigid deformations of the organoid can occur during transportation for microscopy and during the staining procedure, which causes the imaging modalities to not be exactly spatially matched. Future work will aim to develop automatic co-registration of these imaging modalities using deformable 3D image registration techniques [33]. Alternatively, contrast labels that are visible both on MRI and in microscopy could be administered locally to aid in spatial matching.

### 2.2 High-resolution MRI with 3D echo-planar imaging readout

A key challenge for high-resolution microstructural MRI of living organoids is obtaining adequate SNR within acceptable experiment times. In addition to using a 28.2 *T* MRI with a highly sensitive RF-detector, a fast MRI image readout scheme SE–EPI (multiple k-space lines after each excitation) [34] is benchmarked against a conventional SE–linescan (one k-space line per excitation) acquisition. However, EPI is more prone to undesired geometric image distortions due to magnetic field inhomogeneities and variations, caused by differences in tissue magnetic susceptibility and application of diffusion-encoding gradients, which may be exacerbated at high magnetic field and with the use of strong gradients.

Figure 3 compares a dMRI SE acquisition using EPI readout against a linescan readout for dMRI on a phantom sample consisting of two glass capillaries filled with *H*_2_*O* contained 6.3 *mM CuSO*_4_ for shortening measurement time (Figure 3 (a, c)), and a sample with a fixed organoid in phosphate buffered saline (PBS) (Figure 3 (b, d)). Intensity plots across the yellow line (Figure 3 (a, c)) show that the image intensity changes drastically at the interface of glass and water, which indicates that the EPI readout achieves a similar image intensity profile as the linescan readout. Partial volume effects might cause a fraction of a voxel to be filled with water and a fraction with glass, leading to intermediate intensity data points in (Figure 3 (a) and (c)). Additionally, the SE–EPI image of a fixed organoid in PBS (Figure 3 (b)) shows a similar structure and image quality to the SE–linescan readout (Figure 3 (d)).

**Figure 3:**
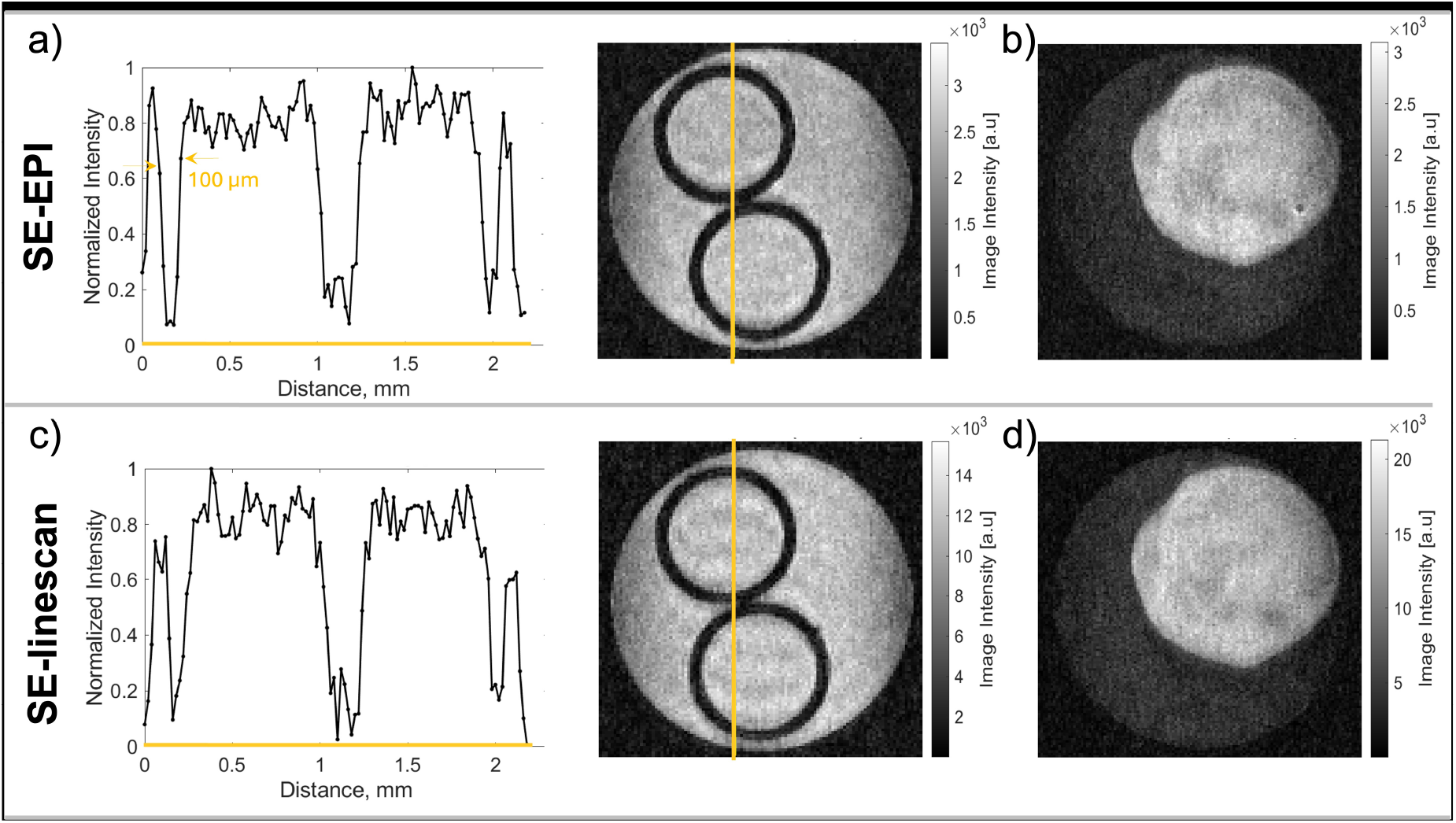
dMRI SE acquisition (using trapezoidal waveforms with *δ* = 0.7 *ms*, Δ = 2.53 *ms* and b-value = 0.5 *ms/µm*^2^) with EPI readout (a, b) and linescan readout (c, d) on a phantom sample and a fixed organoid in phosphate buffered saline (PBS). The phantom consisted of two glass capillaries in *H*_2_*O* with 6.3 *mM CuSO*_4_. The wall thickness of the glass capillaries is 100 *µm*. Panels (a) and (c) display the corresponding normalized intensity profiles along the yellow line for each readout method.

Most previous MRI studies on organoids have relied on conventional linescan readout methods [13, 16, 17, 21–24, 32]. Fast image readout becomes increasingly necessary as field strength increases, because the typically longer *T*_1_ values at higher field strengths result in hours-long experiments to achieve high-resolution MRI. At the 28.2 *T*, the estimates of averaged *T*_1_ and *T*_2_ as measured with SE sequences (RAREVTR and MSME) for the fixed organoid sample are 2000 and 14.8 *ms* respectively, and for a living organoid are 3500 and 14.5 *ms*. Comparison of Figure 3 (b) and (d) shows that the SNR of the SE–EPI experiment with two averages and with a spatial resolution of (20 *µm*)^3^ on a fixed organoid was 8.5 in 4 hours, while the SNR of the SE–linescan was 12.7 in 36 hours. This result indicates that a higher SNR/unit time can be reached using SE–EPI over SE–linescan acquisition.

When performing MRI at voxel sizes on the order of (20 *µm*)^3^ (Figure 3), the actual image resolution can be lower than the nominal spatial resolution set by the experimental acquisition parameters. These spatial resolution limits are known at high-resolution MRI and can be caused by 1) an SNR limitation, 2) line broadening due to diffusion or *T*_2_* relaxation [35, 36]. For the latter factor, high gradient strengths are essential to approach the nominal spatial resolution of the experiment.

At the 28.2 *T* MRI system (3 *T/m* per axis with a slew rate of 30, 000 *T/m/s*, a sufficient SNR was achieved for the spatial resolution of (20 *µm*)^3^.

The limits on spatial resolution can be assessed through the point-spread function, which has shown to have advantages for phase-encoding methods at high spatial resolutions [37, 38]. In this work, the spatial resolution limits were estimated based on a recent study where experimentally derived diffusion limits were reported at different D, *T*_2_* and gradient values for both frequency- and phase-encoding for FLASH-based sequences [39].

The estimation of the resolution broadening in the frequency-encoding direction was done considering *G*_*max*_ = 2 *T/m* (the highest gradient value in [39]), assumed *T*_2_* values in the ranged of 1–10 *ms* and *D* values – 0.5–2 *µm*^2^*/ms* predicted for 28.2 *T* for organoids. Estimating using these values, the actual resolution is 150 percent for nominal resolutions between 3.6 *µm* and 12.2 *µm* which is below the highest used spatial resolution (20 *µm*)^3^ obtained for SE–EPI with frequency- and phase-encoding directions with the gradient strength of 3 *T/m*.

For high-resolution EPI readout, diffusion limits to the image resolution have a more pronounced contribution along the phase encoding direction [40]. In future work, these resolution limits will be further quantified using simulations on microstructural tissue and their experimental validation, specifically regarding high-resolution dMRI sequences with EPI readout.

### 2.3 Microstructural MRI of brain organoids

This Section will describe a series of MRI experiments and associated analyses that probe different aspects of tissue microstructure through diffusion and relaxation, showcasing the versatility of the implemented framework. These measurements overall increase in complexity by increasing the dimensionality of MRI acquisition parameters that are varied (*b*-value, PGSE pulse-separation Δ, gradient direction **g**, diffusion-encoding waveform **g**(*t*), echo time *TE*, and repetition time *TR*). First, changes in organoid microstructure are assessed longitudinally during extended measurement times by the apparent diffusion coefficient (ADC) and volume (Section 2.3.1). Section 2.3.2 probes the signal at higher *b*-values to quantify kurtosis, whereas Section 2.3.3 measures diffusion time-dependent ADC, as signatures of structural complexity. Section 2.3.4 assesses macroscopic anisotropy through estimating the diffusion tensor. Finally, Section 2.3.5 shows high-dimensional correlation experiments in which *b*, **g**(*t*), *TE*, and *TR* are simultaneously varied. The results are discussed in light of corresponding LS microscopy, microstructural characterization of organoids in previous work, and earlier studies in animal specimens.

#### 2.3.1 Longitudinal changes in brain organoids during extended MRI experiments

A particular challenge associated with imaging living organoids is to monitor and ideally prevent undesired structural changes over extended measurement times. As markers for structural changes, the ADC and organoid volume are assessed.

For sufficiently small *δ*, ADC reflects the second cumulant (variance) of the probability distribution function of molecular displacements [41]. As this variance encodes how rapidly molecules have spread over the observed diffusion time, ADC is sensitive to barriers that modulate that spreading such as organelles and cellular membranes. ADC can be estimated from dMRI experiments with a range of b-values where the signal decay is still mono-exponential (typically *b* ≤ 1.2 *ms/µm*^2^ in *in vivo* human brain) for a given diffusion time and gradient direction.

Figure 4 shows ADC- and volume estimates as a function of measurement time for two young and one mature cortical organoids placed in 3 and 5 *mm* glass tubes (Table (1)). In Figure 4 (b), ADC values of all organoids appear relatively constant over time up to 20 hours, with the young 02 and mature 01 organoids showing minimal ADC changes for the entire measurement duration. Likewise, organoid volume over time changes minimally in all organoids until 20 hours after the start of the MRI measurement (Figure 4 (c)). Notably, the volume of the young 02 organoid starts to increase at 23 hours, indicating swelling, as can also be seen in the MRI image insets. This change was accompanied by a decrease and subsequent increase in ADC. Potentially, the young 02 organoid imaged in a tube with an outer diameter of 3 *mm* suffered more from nutrient deficiency than other organoids (young 03 and mature 01), which were placed in 5 *mm* tubes, as a larger volume of culturing medium is surrounding the organoid.

**Figure 4:**
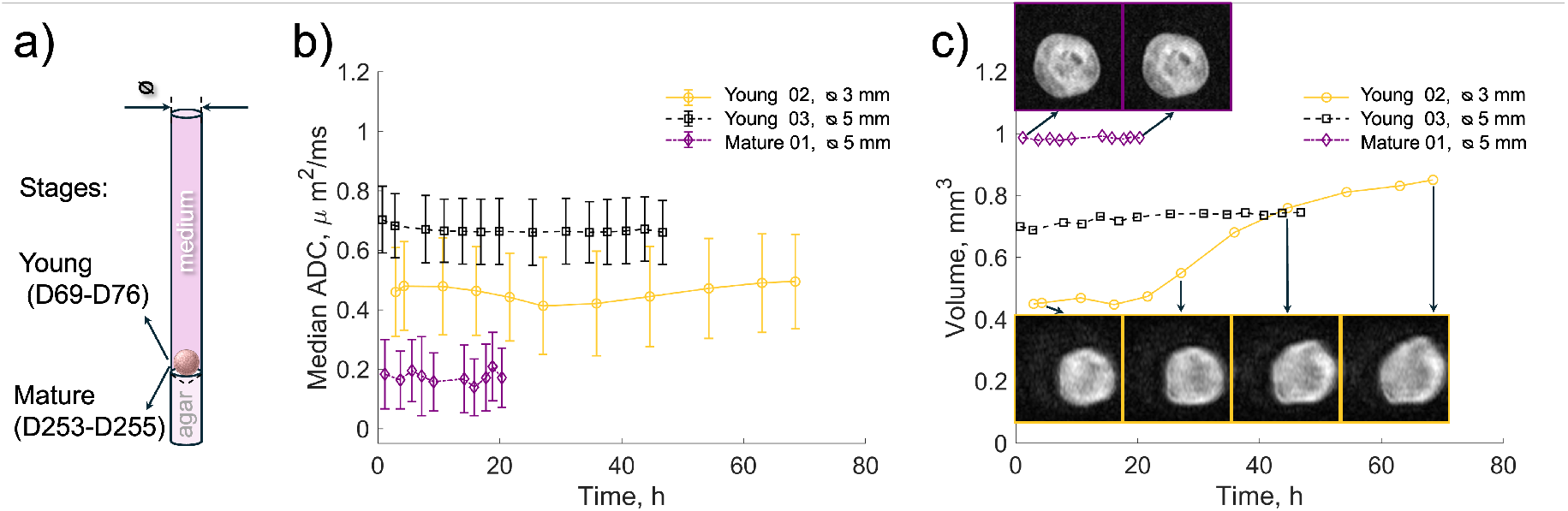
(a) A schematic picture of the MRI tube with a brain organoid atop an agar block in medium. (b) ADC values as a function of measurement time for two young (02 - yellow, 03 - black) and one mature (01 - purple) organoids imaged in 3 and 5 *mm* tubes. The dots represent the median, and the error bars represent the interquartile range across the organoid. (c) Organoid volume for the same three organoids as in (b).

Despite using a high field strength of 28.2 *T* to improve SNR, achieving tens-of-micrometer resolution in dMRI measurements on living organoids necessitates long acquisition times, on the order of hours. Hence, assessing changes in organoid microstructure is crucial for probing living organoids. In this work, organoid structural integrity was inferred from ADC values over time, assuming that a constant ADC value indicates minimal changes in organoid microstructure. The major advantage of this MRI-based evaluation of structural changes is its capacity for longitudinal monitoring, allowing evaluation at the start and end of each MRI scan. In contrast, other viability tests using e.g., staining for apoptosis are destructive and hence only suitable for end-point analyses. Previous work assessed viability by comparing blood gas analysis on medium from two groups of living cortical organoids that did and did not undergo MRI in a 1.5 *ml* Eppendorf tube. This experiment showed no significant change in medium composition and the authors concluded that MRI had no specific negative effect on organoid health [24], yet organoid microstructure was not directly assessed. To extend the viability span of organoids in the MRI setup, additional hardware for exchanging medium, as has been developed for spheroid analysis using NMR spectroscopy [42, 43], could be adapted to extend the measurement time for organoids.

#### 2.3.2 Diffusion kurtosis imaging

Kurtosis analysis can characterize the signal beyond the mono-exponential decay regime as a function of *b* used to estimate ADC [44].

For sufficiently small *δ*, the apparent kurtosis is related to the fourth cumulant of the molecular displacement probability distribution function [41], quantifying its deviation from the Gaussian form. This non-Gaussian behavior may arise from different sources depending on the observed diffusion time: 1) microscopic kurtosis due to restrictions and structural disorder typically within individual microenvironments — such as membranes, organelles, axonal caliber variations, and other hindrances that create complex diffusion pathways on length scales comparable to those probed by the measurement, and/or 2) isotropic- and anisotropic kurtosis due to heterogeneity in size and shape between tissue components [45].

Figure 5 shows estimates of ADC and kurtosis in mature (Figure 5 (a, b)) and young (Figure 5 (d, e)) organoids at 310 *K* at 28.2 *T*, Δ = 4.3 *ms*. Spatial structural heterogeneity within the organoid can be observed both in ADC and kurtosis maps, especially for the mature organoid. At the organoid periphery (3-4 voxels, 150 *µm* − rim) showed higher ADC and lower kurtosis than in the core for both young and mature organoids (Figure 5 (d,e)).

**Figure 5:**
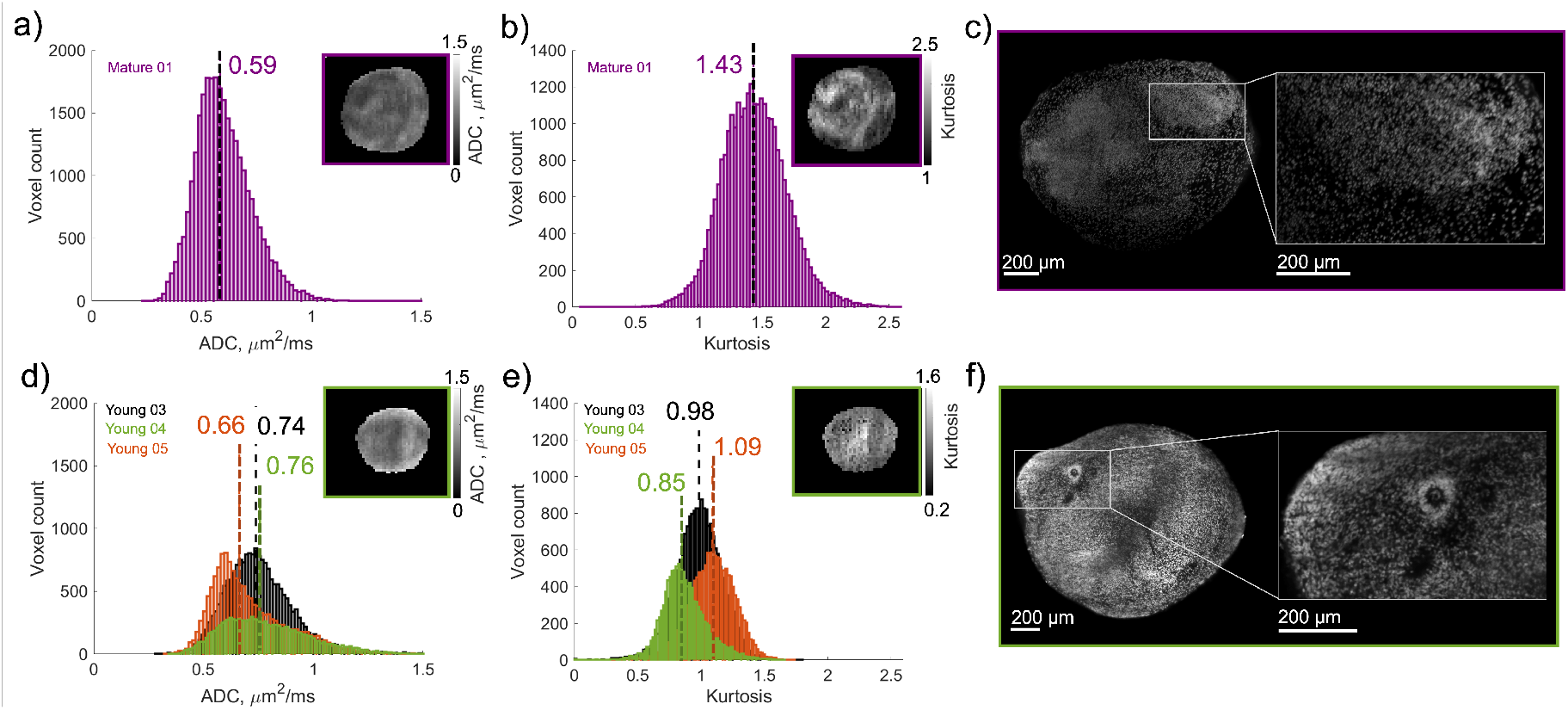
ADC (a, d) and kurtosis (b, e) distributions, depicting values for each pixel within the organoid images, are presented for mature 01 organoid (purple) at the top and young organoids (03 - black, 04 - green, 05-orange) at the bottom. Vertical dashed lines denote median values for each distribution. ADC and kurtosis slice images (maps) are shown in the top right corners of (a, b, d, e) for mature 01 and young 04 organoids, respectively. (c, f) Corresponding LS microscopy images obtained after immunostaining with TO-PRO3 (nuclei).

The analysis of ADC and kurtosis distributions overall showed that median ADC is consistently higher in the younger organoids compared to the mature organoid (median 0.66-0.76 vs 0.55 *µm*^2^*/ms*), while kurtosis is lower (median 0.85-1.09 vs 1.43). Across the younger organoids, ADC and kurtosis distributions vary in median and width. The distributions of kurtosis are narrower in the younger compared to mature organoids.

The spatial heterogeneity of ADC and kurtosis 1) within cortical organoids, and differences 2) across organoids of similar and 3) varying age can originate from different microstructural and experimental aspects. Cortical organoids at any age are spatially heterogenious: the LS images of the same nuclei-stained organoids (Figure 5 (c, f)) show variation of nuclear density within organoids. The spatial heterogeneity can be attributed to local differences in the presence of cell types [29, 46–48], and topographical and cytoarchitectural organization [47]. The variability in ADC and kurtosis values between young organoids of similar age can be explained by differences in their development process. Due to the self-organizing nature of organoids, each individual organoid differs in its cytoarchitecture [29], and organoids exhibit variability in number and size of VLZs. Additionally, organoids are grown in several batches, meaning that their differentiation is started at different timepoints, and derived from different iPSC lines Regarding the observed difference in ADC and kurtosis between young and mature cortical organoids, again developmental changes are likely a key contributor. Nuclear density was overall higher in younger organoids as reported in Section 2.1, which may appear counter-intuitive with higher ADC and lower kurtosis, as higher cellularity (and thus more restriction) is often associated with the opposite trend.

Throughout the development of cortical brain organoids, radial glia and the formation of VLZs are prominent during the early stages [29]. As organoids mature, VLZs disappear and more mature cell populations as astrocytes and mature neurons arise [46, 47, 49]. This increased diversity of neuronal cell types and the evolution of the extracellular matrix from a relatively simple, scaffold-like environment into a more structured, regionally specialized and mechanically functional microenvironment could explain the observed increase in kurtosis in mature brain organoids. Furthermore, the presence and extent of a necrotic core with disintegrated cell membranes due to hypoxia and nutrient deficiency [50, 51] could increase structural disorder. Finally, the variability in microstructural MRI characteristics within and across organoids may be caused by differences in anisotropy at the voxel level (i.e. macroscopic anisotropy, see Section 2.3.4), which affects ADC and kurtosis estimates from MRI experiments with one gradient direction.

Compared to previous work at similarly short diffusion times (3.9 *ms*, Table (1)), the estimated ADC in young organoids here was similar to ADC measured in living rat cortex with oscillating gradient SE (OGSE) (0.73 *µm*^2^*/ms* at an effective diffusion time of 3.8 *ms*, 17.2 *T*, 1 *T/m* gradients) [52]. Yet, the estimated kurtosis in both mature and young cortical brain organoids was higher (≥ 0.59) than in the rat cortex (0.42 at a similar diffusion time). Other work using dedicated strong-gradient human systems and OGSE reported a mean diffusivity of ~1.2 *µm*^2^*/ms* and kurtosis of ~0.75 in cortex, at an effective diffusion time of 4 *ms*, 3 *T*, 200 *mT/m* gradients. Previous work reporting kurtosis measurements in agarose-embedded forebrain-organoids at slightly longer diffusion times using PGSE (Δ = 7 *ms, δ* = 3 *ms*, 11.7 *T*, 1.5 *T/m* gradients, 310 *K*) found notably higher diffusivity and lower kurtosis in the organoid-rim (~0.9 *µm*^2^*/ms* and ~1 respectively) compared to its core (~ 0.6 *µm*^2^*/ms* and 1.5 respectively) [25]. This was attributed to necrotic changes with eosinophilic cell bodies and pyknotic nuclei observed on 2D hematoxylin and eosin staining in the core, and the increased presence of neurites and glial processes in the rim, which was above 600 *µm* based on shown dMRI images [25]. No direct qualitative or quantitative spatial comparison was presented between MRI and microscopy in the same organoid and the age was not specified, but the results describe microstructural features for rather young organoids.

The differences compared to previous studies may be explained by differences in organoid types and their microstructure, organoid embedding (agarose vs medium), and hardware. Differences in field strength cause different compartmental apparent relaxation times and, as such, a different weighting of compartments (intra- and extra-cellular) in the signal.

#### 2.3.3 ADC diffusion-time dependence

The ADC can be probed as a function of diffusion time (here achieved by varying Δ), and its behavior reflects characteristics of the microstructure. For very short times molecules only sense their immediate neighborhood per the local intrinsic diffusivity of the medium; for longer times coarse graining occurs modulated by structural disorder and exchange; and finally ADC reaches its long-time limit when full coarse graining within each interconnected tissue compartment has taken place [2, 53]. The MRI gradient hardware used here can enable measurement of much shorter diffusion times than with typical clinical scanners.

Figure 6 illustrates the impact of varying Δ on the dMRI signal and ADC in different regions of the young and mature cortical organoid. Across the organoid, signal intensity visibly increases with increasing Δ in both organoids (top row of (a) and (c)). Concomitantly, ADC overall decreases as a function of diffusion time in most regions, yet the diffusion-time dependence is markedly different in the young and mature organoids. Whereas the older organoid shows the steepest change in ADC between Δ = [2.7, 10] *ms* both globally and regionally, the younger organoid shows a slight ADC increase followed by a decrease in the periphery (p = 5.5e-52 (mask 4) and 6.4e-99 (mask 5) between Δ = 2.7 and 5 *ms*; and p = 7.2e-12 (mask 4) and 2.371e-13 (mask 5) between Δ = 2.7 and 10 *ms*), and a more gradual linear diffusion-time dependence in core regions across the whole range of measured diffusion times Δ = [2.7, 25] *ms*. Moreover, similar to Figure 5, a higher ADC is observed in the younger organoid, and this trend holds for all diffusion times.

**Figure 6:**
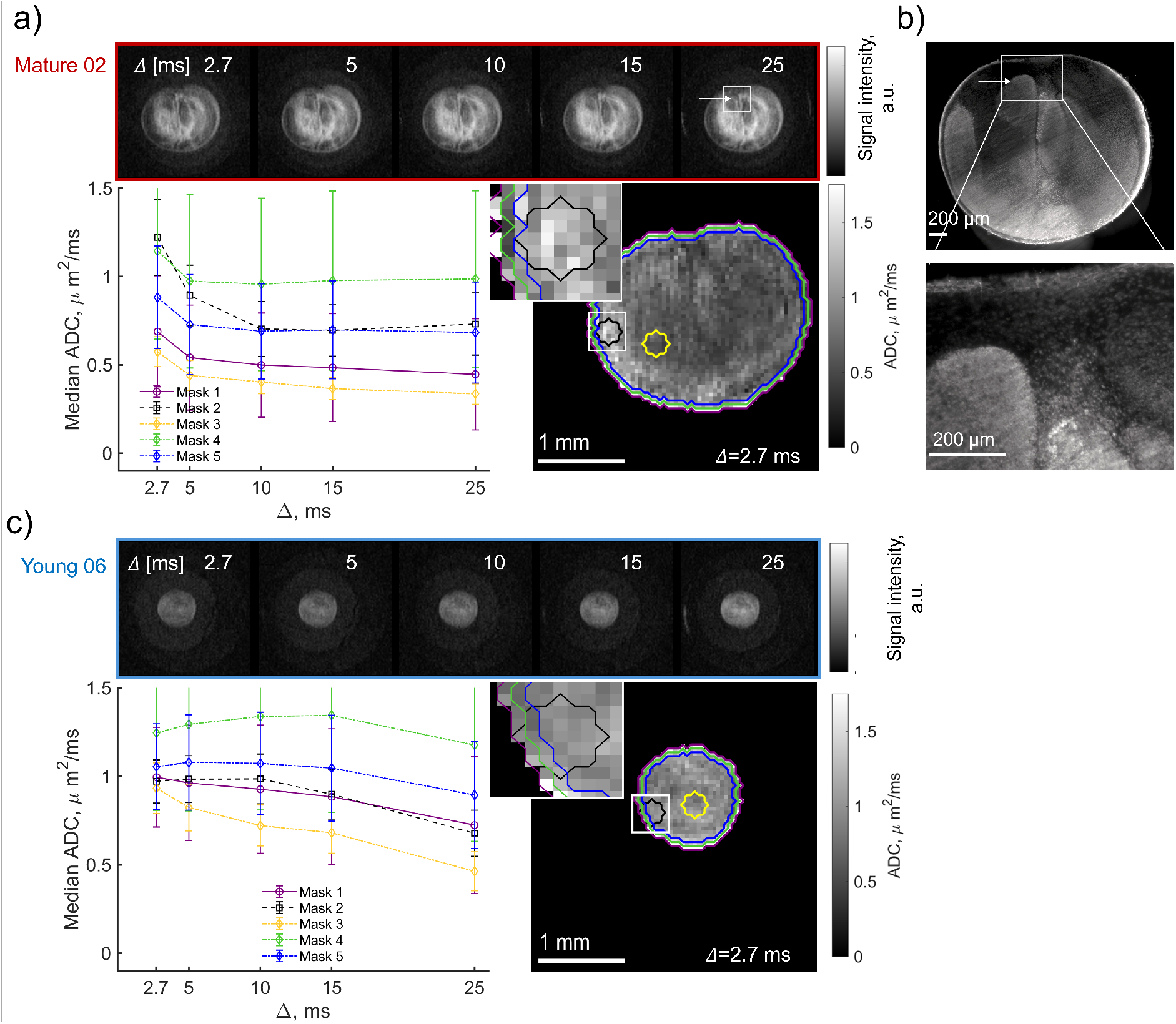
The diffusion time-dependence of ADC is illustrated for mature 02 (red) in (a, b) and young 06 (blue) (c) organoids. (a) For the mature 02 organoid, the dMRI signal intensity maps at *b* = 1 *ms/µm*^2^ as a function of Δ are shown (top row). The median ADC is calculated for the entire organoid (1-purple), as well as two smaller regions 2-black, 3-yellow) and two after each other circumferences at the edge of the organoids selecting a ring with one voxel thickness of 44.8 *µm* (4-green, 5-blue) (bottom-left). The results are plotted as a function of Δ to assess diffusion-time dependence of ADC. The error bars show standard deviation of the ADC values in selected regions. The ADC maps at *b* = 1 *ms/µm*^2^ and Δ = 2.7 *ms* are displayed alongside (bottom-right). (b) LS microscopy images obtained after immunostaining with TO-PRO3 (nuclei) for the mature 02 organoid. (c) The young organoid underwent the same data analysis as the mature organoid in (a).

The comparison of MRI with LS microscopy images with nuclear staining (Figure 6 (a, b)) was performed for the mature organoid 02 and revealed areas of similar spatial features in nuclear density distributions and dMRI signal intensity. The area indicated by the white box and arrow shows lower dMRI signal intensity where nuclear density appears higher. Compared to dMRI signal intensity, the spatial variability in ADC is visibly less pronounced, suggesting contributions from differences in *T*_2_ across the organoid that effect both the non-diffusion weighted and diffusion-weighted images. Yet quantitatively, the estimated ADC in the periphery (rims of one voxel in width (Figure 6 (a), masks 4, 5) and a region of 300 *µm* − thickness (Figure 6 (a), mask 2)), are higher than in the center of the organoid (mask 3). This is consistent with the organoids studied in Section 2.3.2 and previous work [25]. For the young organoid (Figure 6 (c)), the dMRI results showed again higher ADC values at the periphery regions in comparison to regions near the core of the organoid. This organoid was lost during the staining procedure which did not allow for a direct comparison of nuclear density distribution.

In fixed forebrain organoids, previous work measured both time-dependent mean diffusivity and kurtosis [25], providing microstructural markers related to effective cell size and membrane permeability.

dMRI experiments with both short (Δ = 7 *ms, δ* = 3 *ms*) and an order-of-magnitude longer (Δ = 200 *ms, δ* = 3 *ms* with stimulated echoes (STE) to circumvent the significant signal decay from *T*_2_-relaxation) diffusion times were used. In the shorter diffusion-time regime, a peak in the time-dependent kurtosis curve *t*_peak_ was observed and explained by the occurrence of two competing phenomena: on the one hand the restriction of water diffusion by barriers and the associated increase of kurtosis, and on the other hand the exchange of water across permeable membranes and the associated decrease of kurtosis [52, 54, 55]. Characterizing *t*_peak_ can thus provide information on membrane permeability, intrinsic compartmental diffusivities, cell size, and cell density. The authors show that at body temperature (310 *K*), *t*_peak_ was shifted beyond the shortest measured diffusion time, with generally even shorter *t*_peak_ values in the rim (with a width ≥ 600 *µm*) compared to the core [25]. The hardware setup used in the current work enables shorter diffusion times and thus further characterisation of *t*_peak_ in future work. This can be further extended to oscillating gradients for even shorter diffusion times to estimate surface-to-volume ratio and intrinsic diffusivity [56]. In the long diffusion-time regime, the effect of exchange dominates and the diffusivity becomes constant as a function of time (complete coarse graining), whereas the time dependency in kurtosis can be used to estimate exchange time [44, 57]. To achieve long diffusion times at ultra-high field strengths, STE implementation is necessary because of the even shorter apparent *T*_2_, and future work will develop robust STE acquisitions at a coarser image resolution, ensuring the diffusion length to remain within the voxel dimensions. Mean diffusivity was found to decrease as would be expected from coarse graining. Increases in ADC – as observed here in the rim of the young organoid – have been observed in other studies in gray matter, albeit at longer diffusion times ≥ 16 *ms* [58]. This could possibly be attributed to kurtosis effects biasing ADC estimates at the *b*-values used and requires further investigation.

#### 2.3.4 Diffusion tensor imaging

Macroscopic fractional anisotropy (macro-FA), which is for example high in voxels with coherently oriented neurites or cells (e.g. in VLZs), can be characterised by measuring a series of images probing diffusion in different directions and estimating the diffusion tensor - a 3-dimensional extension of ADC.

Figure 7(a, c) presents the mean diffusivity (MD, mean of the tensor eigenvalues) in grayscale and combined color-coded mean diffusion direction with color brightness modulated by macro-FA (cFA) for two young organoids, 07 and 08, at the age of D70 and D73, respectively. For both organoids, the MD values were mainly in the range 0.4 − 0.9 *µm*^2^*/ms* and were similar to previous reports [25]. Regions of high MD (0.75 − 0.9*µm*^2^*/ms*) and macroscopic cFA (0.5 − 0.75) around VLZs resemble the radial alignment that is typical for the developing cerebral cortex [46, 48, 49]. For organoid 08, VLZs were not clearly visible likely due to lower spatial resolution ((30 *µm*)^3^ for organoid 08, and (20 *µm*)^3^ for organoid 07). Figure 7(b, d) shows corresponding LS microscopy images showing VLZs (green), nuclei (pink), and axons (yellow). Comparing MRI and LS microscopy, the VLZs show a higher MD despite the higher nuclear density (Figure 7(a, b)). The organoid 08 LS images (Figure 7(d)) showed that VLZs were smaller in size compared to organoid 07, which can explain why they are difficult to disentangle on MRI (Figure 7(c)). As also observed in the ADC images in Sections 2.3.2 and 2.3.3, regions of higher MD correspond to regions of high nuclear density (e.g. around the VLZs), which appears counter-intuitive as MD is expected to be inversely proportional to cell density with higher cell density restricting water movement. Nevertheless, also FA is higher around the VLZs, indicating less-restricted diffusion in at least one direction which can result in an overall higher MD.

**Figure 7:**
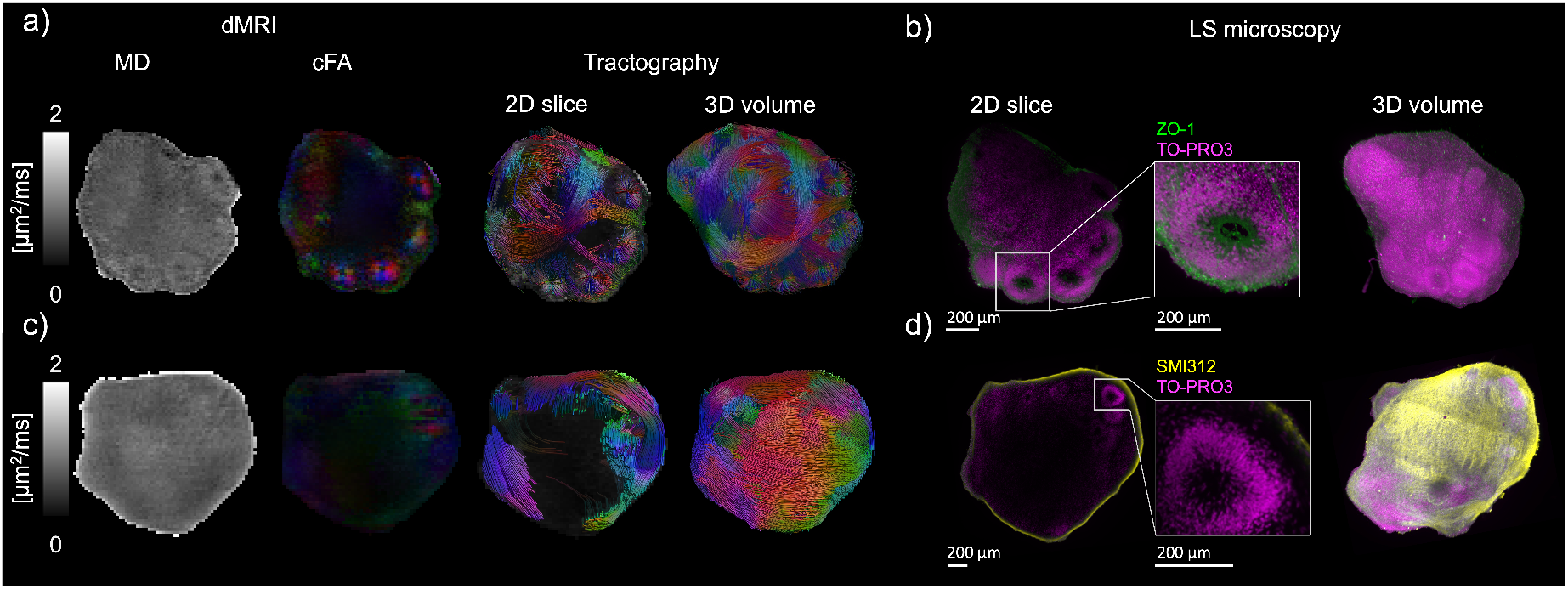
Diffusion tensor imaging metrics are presented alongside structural information obtained from lightsheet (LS) microscopy for two young organoids, 07 and 08. Specifically, (a, c) show mean diffusivity (MD), color-coded macroscopic fractional anisotropy (cFA). The cFA shows the color-coded primary diffusion direction (red for left-right, blue for top-bottom, green for parallel to the viewing axis) with color brightness modulated by FA. Additionally, tractography reconstructions by seeding streamlines from the visualised 2D slice and from the entire 3D volume are presented. (b, d) 2D and 3D LS microscopy images for approximately the same slice and orientation as MRI. The colors of the LS microscopy image correspond to the following: green channel − ZO-1 (VLZs), yellow channel − SMI312 (axons), and pink channel − TO-PRO3 (nuclei).

Our results show the potential of MRI as alternative non-invasive 3D characterization of living cortical brain organoids to investigate cell density patterns and structures like VLZs. Intact organoid imaging with microscopy has only relatively recently resulted in new insights of microstructure, e.g. cell density distributions in VLZs and cortical plate, and alternations in disease [59]. In addition, VLZ count and size can provide information on the proliferation of radial glial cells, which may be altered in organoids modelling disease [60]. In previous work, the VLZ-structure was investigated in bioprinted cortical organoids on a 3T system [23] and visualized by weighting the MRI signal by the *T*_1_ relaxation time. Hyperintense signals appeared to correlate with areas of higher cellularity. Other work has manually traced VLZs on *T*_2_-weighted images [21]. The current work showcases an extensive microstructural MRI framework relying on diffusion-weighting as a physical phenomenon direct marker modulated by microstructure, whereas *T*_1_ can be regarded as more indirect marker through its sensitivity to water- and macromolecular content. The showcased framework here enables combining multiple contrasts including diffusion, *T*_1_ and *T*_2_ for detailed assessment that integrates the complementary information from diffusion and relaxation MRI (see Section 2.3.5).

Tractography reconstructions (Figure 7(a, c)) enabled the visualisation of discrete streamlines that cover predominantly the periphery of the organoid, but also traverse around and through the VLZs thereby exposing their radial organization. The LS microscopy images obtained for axon staining (Figure 7(c)) confirm that high-macroscopic FA regions and reconstructed tractography streamlines correspond to fiber-like structures that form a network organization in organoids. Tensor- and tractography information can aid co-registration between MRI and microscopy of the same specimen based on microstructural features [33, 61, 62]. One previous study performed tractography on organoids at 11.7 *T*, where organoids were fixed and a contrast agent was used to increase the SNR/unit time [21]. This enabled the visualisation of longer streamlines in the organoid periphery and at the perimeter of the circular protrusion, and interconnections between VLZs. In the current study, we showed the feasibility of diffusion tensor imaging and subsequent tractography analysis in living organoids in culturing medium at high resolution, without the addition of a contrast agent. Moreover, with the over 3-fold smaller voxels, also fiber-like structures within the VLZs could be visualised.

#### 2.3.5 Frequency-dependent diffusion tensor-relaxation distributions *D*(*ω*)-*R*_1_-*R*_2_

While the preceding sections focused on conventional trapezoidal diffusion waveforms that each probe diffusion along a single axis and at a single diffusion time per measurement, additional and complementary information can be accessed by moving beyond this framework. By implementing more complex gradient waveforms that yield *b*-tensors beyond rank-1 (Figure 1 (b)) and the joint integration of longitudinal *R*_1_ = 1*/T*_1_ and transversal *R*_2_ = 1*/T*_2_ relaxation contrasts, microstructural heterogeneity at a sub-voxel level can be better disentangled compared to standard acquisitions. This approach was suggested for obtaining extensive information on the diffusion time-dependency in biological tissues via a unified, (biophysical) model-free framework [63–65]. To this end we adopt an approach that represents the MRI signal in each voxel by a distribution of diffusion tensors with varying fraction, shape, size, orientation, frequency (*ω*) dependence, and relaxation properties **D**(*ω*)-*R*_1_-*R*_2_ [66, 67]. The estimation of such a joint diffusion-relaxation tensor distribution requires an extensive acquisition probing a wide range of *b*-values, diffusion frequencies *ω*, encoding *b*-tensors, TE, and TR [64]. A flexible custom MRI sequence was implemented at the 28.2 *T* system that enabled the execution of arbitrary diffusion-encoding waveforms and controlled adaptation of TE and TR (Figure 1 (b)), beyond what is possible with vendor-supplied sequences. The high gradient strength and slew rate of the hardware used in this study allow for sampling a wide range of frequencies from 50 to 300 *Hz* with gradient waveform durations between 3 and 9 *ms*.

The rich multidimensional information of the **D**(*ω*)-*R*_1_-*R*_2_ distributions is condensed into parameter maps of their means *E*, variances *V*, and covariances *C*. Figure 8 shows per-voxel statistical representation of means *E* and variences *V* for isotropic diffusivity *D*_*iso*_, normalized microscopic diffusion anisotropy *D*_Δ_^2^, *R*_1_, *R*_2_, and frequency-dependence (Δ_*ω/*2*π*_), assessed at *ω/*2*π* = 51 *Hz*. The means *E*[*D*_*iso*_], *E*[*R*_1_], and *E*[*R*_2_] correspond most closely to conventional mean diffusivities (Figures 5, 6, 7), *R*_1_ and *R*_2_, respectively. Analysis of the relaxation properties of the cortical organoid showed that means *E*[*R*_1_] (Figure 8 (a) varied between 0.1 − 0.37 *s*^−1^ and was spatially homogeneous, whereas means *E*[*R*_2_] was in the range 25 − 65 *s*^−1^, with lower values in the center of organoid. The *E*[*D*_Δ_^2^] maps (Figure 8 (a, c)) quantify microscopic diffusion anisotropy which, if the tensors are coherently oriented at the voxel level, leads also to a high macroscopic anisotropy that can be captured with conventional diffusion tensor analysis (Figures 7). Higher *E*[*D*_Δ_^2^] can be observed near the periphery, corresponding to the presence of VLZs (Figure 8 (d)) and axons (Figure 7). The frequency dependence maps (Figure 8 (b)) show a rate of change with frequency within the explored range of 50 − 300 *Hz*. Positive values of the Δ_*ω/*2*π*_*E*[*D*_iso_] are characteristics of restricted diffusion [63, 64, 68], while negative values in Δ_*ω/*2*π*_*E*[*D*_Δ_^2^] indicate a decrease in anisotropy with higher frequency at the short diffusion time regime. As diffusion frequency is inversely related to diffusion time, the obtained results are coherent with the decrease of ADC in Figure 6. The non-zero variances V (Figure 8 (a, b)) indicate the intravoxel heterogeneity and highlight voxels comprising multiple water populations with different diffusion and/or relaxation properties [63, 64]. Additionally, the increase in variances can be caused by contributions from non-Gaussian diffusion effects and include microscopic kurtosis [63], which is not explicitly disentangled in the **D**(*ω*)-*R*_1_-*R*_2_ [64]. Variances were overall higher in the organoid periphery, and for *D*_iso_ and *R*_2_ also in the organoid core (Figure 8 (a, b)).

**Figure 8:**
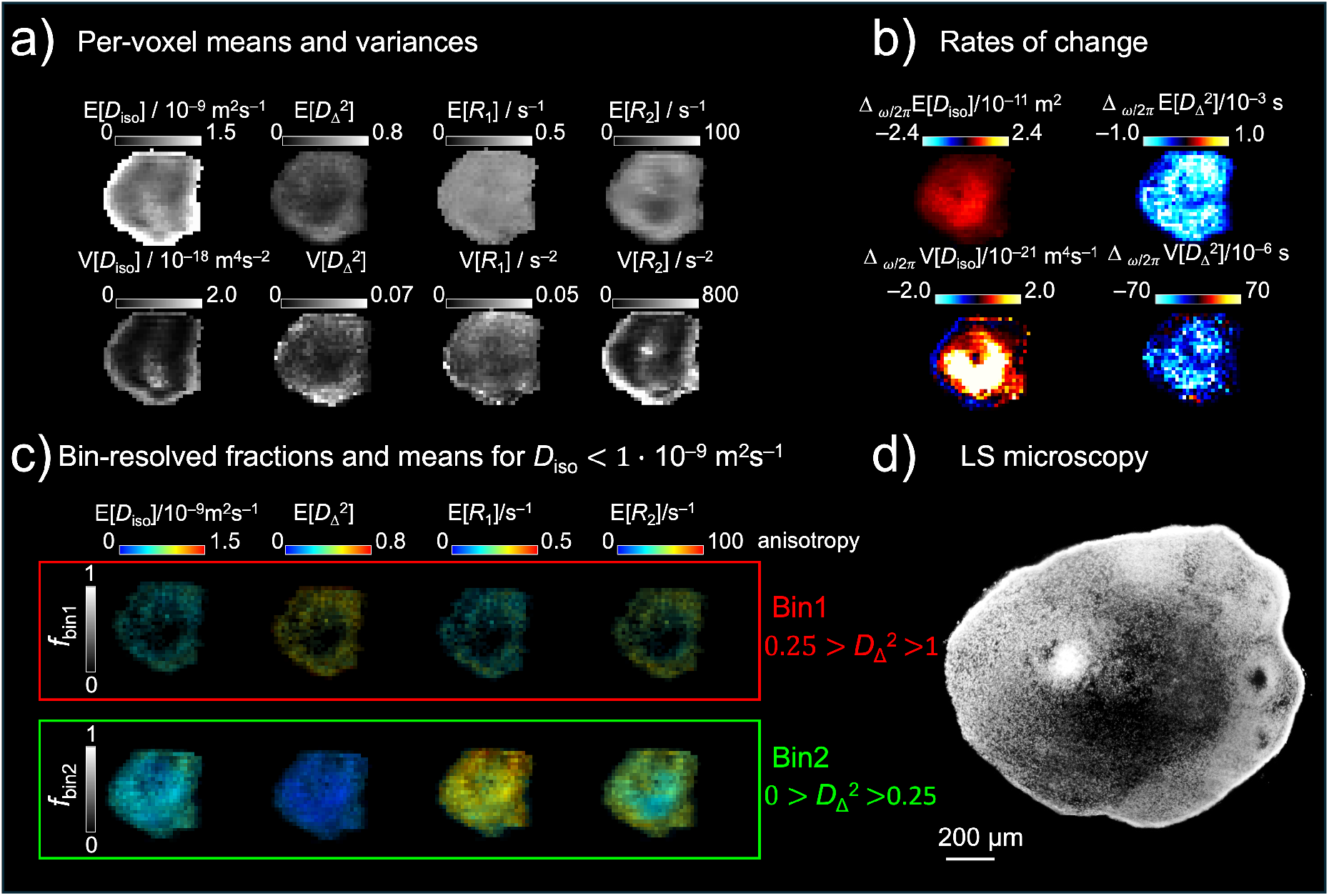
Parameter maps derived from the per-voxel *D*(*ω*) − *R*_1_ − *R*_2_ distributions of the *in vivo* young cortical organoid 07, slice 40. (a) Per-voxel means *E* and variances *V* of the frequency dependent isotropic diffusivity *D*_*iso*_, normalized diffusion anisotropy *D*_Δ_^2^, longitudinal *R*_1_ and transversal *R*_2_ relaxation rates. (b) Frequency dependence (Δ_*ω/*2*π*_) through per-voxel rates of change of means *E* and variances *V* for *D*_*iso*_ and *D*_Δ_^2^ calculated at the low sampling frequency of *ω* = 51 Hz and their corresponding Δ_*ω/*2*π*_ values between *ω* = 51 and 214 Hz. (c) Vizualization of compartments with low diffusivity and high/ low anisotropy for means *E* and variances *V* for *D*_*iso*_, *D*_Δ_^2^, *R*_1_ and *R*_2_. The bin limits are selected at the low *ω* value of 51 *Hz*. (e) LS microscopy image obtained after immunostaining with TO-PRO3 (nuclei) for young organoid 07.

For further visualization, the resolved distribution components were manually clustered into bins. For the cortical organoids, two bins were defined corresponding to classes with anisotropic and lower diffusivity tensors (bin 1), and isotropic and lower diffusivity tensors (bin 2) [64]. Most signal components are clustered into bin 2, whereas higher anisotropy tensors (bin 1) can be seen on the periphery of the organoid which is in agreement with high values of means *E*[*D*_Δ_^2^] and can be explained by VLZs and axonal presence (Figure 7). The compartment visualization (Figure 8 (c)) showed that the means *E*[*R*_1_] of the high anisotropy bin 1 were lower than values for the low anisotropy bin 2. This result in line with *R*_1_ values reported for biophysical modelling of soma, extra-neurite and intra-neurite compartments using the Soma And Neurite Density Imaging (SANDI) model [69]. The authors showed that that the intra-neurite *R*_1_ was lower than soma and extra-neurite *R*_1_ at a field strength of 3 *T*. Using the same biophysical model, compartmental *R*_2_ showed a similar trend [70] which is in line with lower *E*[*R*_2_] for the anisotropic bin 1 here in the periphery. The center of the organoid shows an opposing trend in *E*[*R*_2_], likely due to areas of necrosis.

### 2.4 Outlook

This work demonstrates the feasibility of performing detailed and high-resolution microstructural MRI at ultra-high magnetic field on living cortical brain organoids, followed by 3D lightsheet microscopy. A variety of microstructural MRI acquisition strategies, including diffusion encoding with trapezoidal and arbitrary waveforms, were implemented with fast readouts, revealing structural heterogeneity, macro- and microscopic anisotropy, the feasibility of longitudinal monitoring, and differences in microstructure between young and mature brain organoids.

The uniqueness of the 28.2 *T* system – currently the only commercially available system at this field strength equipped for imaging – enables high SNR and thus high spatial resolution microstructural MRI up to (20 *µm*)^3^ within acceptable scan times, making it feasible to sample a wide range of diffusion- and relaxation-acquisition parameter settings. At the same time, however, the uniqueness of the system limits the widespread applicability of the measurements implemented here. Yet, it is envisioned that the unique detailed imaging that this system provides can contribute to answering longstanding questions in the field of microstructural MRI [71]. The development of appropriate biophysical models to interpret the MRI signal in terms of relevant tissue microstructural parameters has been an active area of research for the past decades. Inaccurate model assumptions can lead to biased results and interpretations. Examples include the validation of assumptions on intrinsic diffusivities [72, 73], the ability to measure axon and cell soma diameters with MRI [74–76], and quantification of typical exchange times [77–79]]. Furthermore, the proposed MRI platform can be used for extensive protocol optimisation to capture MRI biomarkers with high precision and accuracy. With diffusion being a physical process independent of the MRI field strength, the obtained results can be translated to lower field strengths if relaxation processes and susceptibility effects are appropriately taken into account.

Nevertheless, the sensitivity that ultra-high field provides becomes necessary when high spatial resolutions are required to image increasingly small structures; whereas cortical brain organoids are relatively large, other organoids (e.g. tumoroids) can be much smaller, in the order of 50-1000 *µm*. The current setup thus provides an optimal platform for detailed microstructural investigation of a wider range of organoids with MRI. Further developments in hardware on ultra-high field MRI systems (e.g. microcoils [80] or cryoprobe setups [21]) would allow for higher SNR, whereas stronger gradients would allow for even faster diffusion encoding. Image acquisition strategies, such as compressed sensing and super-resolution reconstruction, can further push the resolution [81–83].

The microstructural MRI toolbox implemented in this work can be extended in several ways. The flexibility of the arbitrary-waveform sequence in principle enables complete freedom in diffusion-encoding strategies, within hardware limits. STE sequences will furthermore enable measurements at long diffusion times in future work. In addition to measuring microstructural information from the diffusion of water molecules, metabolic imaging can provide a new window into investigating the function of organoids. Preliminary results using the setup in this work [30] have shown that while organoid structure appears to remain intact over extended measurement times (Figure 4), the signal intensity of measured lactate increases, indicating potential early changes in metabolism before structural changes occur. Moreover, combining microstructural and metabolic imaging with diffusion magnetic resonance spectroscopy [84] can provide more cell-specific measurements by measuring metabolite diffusion rather than water diffusion, with the latter being ubiquitous in all intra- and extracellular spaces.

Leveraging the full suite of implementable MRI techniques, organoid MRI enables microstructural characterization and longitudinal monitoring of changes in response to pharmacological or environmental interventions in a controlled, human-relevant model system. By bridging MRI biomarkers in organoids and *in vivo*, this framework lays the groundwork for patient-specific translation, ultimately supporting more precise diagnosis, treatment stratification, and therapeutic monitoring in the clinic.

## 3 Methods

### 3.1 Samples

#### 3.1.1 Phantom

The validation phantom consisted of an NMR glass tube with 2.5 *mm* outer diameter (Hilgenberg) in which two glass capillaries with an outer diameter of 1 *mm* and wall thickness 100 *µm* (Bruker) were inserted. The NMR tube and capillaries were filled with a reference solution, which consisted of H_2_O with 6.3 *mM* CuSO_4_ (Sigma-Aldrich).

#### 3.1.2 Organoids

IPSCs were cultured in StemFlex (Life Technologies, A3349401) on Geltrex (Gibco, A1413202) coated dishes. Cells were passaged when 80-90 % confluency was reached, using 0.5 *mM* EDTA (Invitrogen, 15-575-020). All lines were frequently tested for mycoplasma (MycoAlert kit, Lonza Bioscience, LT07-318). Cortical organoids were generated according to Yoon et al. [29], with some adaptations. In short, 9000 cells per EB from lines (929C4 [85], CS29iALS-C9n1 and CS29iALSC9n1.ISOT2RB4 [86]) were seeded in hES0 medium (20 % KOSR (Gibco, 10828028), 3 % FBS (Sigma-Aldrich, F7524), 2 *mM* L-Glutamine (Gibco, 25030024), 1X MEM-NEAA (Gibco, 11140035), 0.0385 *mM* 2-mercaptoethanol (Fisher Scientific, 11441711) in DMEM-F12 (Gibco, 11320074)), supplemented with 50 *µM* Y-27632 (Axon Medchem, 1683) and 4 *ng/mL* FGF2 (PeproTech, 100-18B), in an ultra-low attachment 96-well plate. On day 2 and day 4 the medium was changed to hES0 medium supplemented with 2.5 *µM* Dorsomorphin (R&D Systems, 3093) and 10 *µM* SB-431542 (Axon Medchem, AXON 1661). From day 6 to day 25 the medium was changed three times per week with neural medium (1X B27 without vitamin A (Gibco, 12587010), 2 *mM* L-Glutamine, 100 *U/mL* penicillin-streptomycin (Gibco, 15140122) in Neurobasal medium (Gibco, 21103049)), supplemented with 20 *ng/mL* EGF (R&D Systems, 236-EG) and 20 *ng/mL* FGF2. From day 25 to day 43 the medium was changed three times per week to neural medium with 20 *ng/mL* BDNF (STEMCELL Technologies, 78005.1) and 20 *ng/mL* NT-3 (PreproTech, 450-03B). On day 43, organoids were transferred to 6 *cm* petri dishes. From day 43 until harvesting, the medium is changed three times per week with neural medium without any supplements.

On the day of the MRI experiments, organoids were transferred into MR tubes (Bruker) of either or 5 *mm* outer diameter (Table 1) atop an 2 % agarose substrate (Sigma-Aldrich Cat*#*11388983001). The MR tubes with agarose substrate were prepared one day prior the organoid transfer. The agarose was placed into the MR tubes in liquid form using a pasteur pipette, forming a slight concave meniscus that provided a resting platform for the organoid. After the agarose solidified, the tube was filled with medium and kept at 37 °*C* to stabilize the agarose. Before transferring the organoid, the medium was exchanged with fresh medium by filling completely the MR tube. The organoid was then transferred by pipetting it into the medium surface on top the the MR tube, allowing the organoid to drop and land atop the agarose block.

### 3.2 MRI

A 28.2 *T* NMR spectrometer with an Avance Neo console and ParaVision360 V3.3 was used with an IProbe containing a 5 *mm* volume coil and a gradient insert of 3 *T/m* with a slew rate of 30,000 *T/m/s* (Bruker, Biospin GmbH, Germany). The probe temperature was set to 37 °*C*.

Shimming was performed on each organoid at the beginning of the MRI measurements to optimize the magnetic field homogeneity. After temperature stabilization and optimization of transmitter frequency, first-order shims were adjusted, followed by an 11-step FID shimming procedure that included first-order, second-order, and z^3^ shims. Alternatively, a so-called Mapshim protocol was used. The shimming protocols were optimized for each specimen.

*T*_1_ and *T*_2_ measurements were performed to estimate the contribution of the relaxation and estimate repetition time (TR) and therefore experimental times. For *T*_1_, a fast spin echo sequence with variable repetition times (RAREVTR) was used with the following parameters for a fixed organoid sample placed in 3 *mm* tube: TR [300-3000] *ms*, RARE factor 1, echo time (TE) 2.79 *ms*, matrix size 96 x 96, field of view (FOV) 2.6 x 2.6 *mm*^2^, slice thickness of 100 *µm*, 1 average, receiver bandwidth 100,000 *Hz*. For *T*_2_ determination, a spin echo sequence was used (MSME) with the following parameters, respectively, for the fixed organoid sample: TR 8000 *ms*, TE 3.5 *ms*, matrix size 96 x 96, FOV 2.6 x 2.6 *mm*^2^, slice thickness of 100 *µm*, 4 averages, receiver bandwidth 100,000 *Hz*. Note that the *T*_2_-values are determined from spin-echo images at high spatial resolutions and therefore the observerd *T*_2_-values might deviate from the intrinsic *T*_2_ values [87, 88].

Microstructural MRI measurements with diffusion encoding were performed with a spin echo (SE) sequence and 3D echo-planar imaging (EPI) readout with diffusion-encoding gradients **g**(*t*) that were either trapezoidal or arbitrary in shape (Figure 1 (b)). The axially symmetric positive semi-definite b-tensor characterising the diffusion encoding was computed as 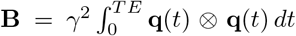, where 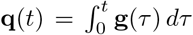 and *γ* is the gyromagnetic ratio. The *b*-value indicating the strength of diffusion encoding was derived as *b* = Tr(**B**), where Tr is trace, and the main diffusion encoding direction as the first eigenvector of **B**. The anisotropy of the b-tensor was characterised by *b*_Δ_ = (*b*_∥_ − *b*_⊥_)*/b*, where *b*_∥_ and *b*_⊥_ are the eigenvalues corresponding to the eigenvectors along and perpendicular to the symmetry axis, respectively [65, 89, 90]. For trapezoidal waveforms (also called pulsed-gradients (PG)), *b* = −*γ*^2^*g*^2^*δ*^2^(Δ − *δ/*3) with gradient-lobe duration *δ*, gradient-lobe separation Δ, and strength *g* (Figure 1 (b), left), and *b*_Δ_ = 1.

First, the evaluation of image quality of dMRI with 3D EPI readout vs conventional linescan image readout was investigated. Next, quantitative microstructural characterization in living organoids with a range of dMRI experiments was performed.

#### 3.2.1 High resolution dMRI with 3D echo-planar imaging readout

PGSE experiments (Figure 1 (b)) were evaluated using EPI readout against a conventional linescan readout in a phantom and fixed organoid). For the phantom, the following parameters were used: TR = 1000 *ms*, TE = 7.49 *ms*, FOV (2.2 *mm*)^3^, matrix (100)^3^, spatial resolution (20 *µm*)^3^, *δ* = 0.7 *ms*, Δ = 2.53 *ms*, 1 diffusion encoding direction, b-value = 0.5 *ms/µm*^2^, 4 averages. The EPI readout had a receiver bandwidth 277,777 *Hz*, 8 segments, total experiment time 1 *h* 57 *min*. The linescan readout had a receiver bandwidth of 100,000 *Hz* and a total experiment time 15 *h* 38 *min*. A signal intensity analysis was performed using a home-written Matlab script (Matlab R2024b).

For the fixed organoid, the following parameters were used: TR = 3500 ms, TE = 9.1 ms, FOV (2.6 *mm*)^3^, matrix (128)^3^, spatial resolution (20 *µm*)^3^, *δ* = 1 *ms*, Δ = 4 *ms*, 1 diffusion encoding direction, b-value = 1.0 *ms/µm*^2^, 2 averages. EPI readout had receiver bandwidth 322,580 *Hz*, 8 segments, total experiment time 3 *h* 59 *min*. The linescan readout had receiver bandwidth 100,000 *Hz* and total experiment time 36 *h*.

SNR calculations were computed in MATLAB R2024a as the mean foreground signal divided by the mean background signal multiplied by 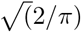 (to correct the noise-standard deviation estimate for Rician bias) [91].

#### 3.2.2 Microstructural assessment of cortical brain organoids with dMRI

A wide variety of microstructural MRI acquisitions on living cortical organoids were performed with varying gradient waveforms, b-values, diffusion-times and frequencies, and gradient directions (see Table 1 for an overview), described in the next paragraphs. For all dMRI data, binary image masks were created using a custom MATLAB script (MATLAB R2024b), employing thresholding and edge detection on a dMRI image (typically *b* ~ 1 *ms/µm*^2^ was found to have a good contrast), followed by erosion to remove one to two voxels from the image edges. This erosion step was implemented to mitigate partial volume effects with the background. dMRI data were denoised using tensor MPPCA [92]. Additionally, the diffusion tensor imaging data were corrected for eddy current geometric distortions in FSL [93, 94]. Note that not all preprocessing steps were appropriate for all datasets because of the number of images and distribution of gradient directions.

##### Longitudinal changes in organoids during extended MRI experiments

Three organoids underwent repeated longitudinal dMRI (Table 1, longitudinal experiment). The effect of tube size (3 vs 5 *mm* diameter) was investigated on two young 01(D69) and 02(D74), and one mature 01 (D253) organoids. Shimming was performed at the start of the longitudinal experiment.

ADC can be estimated from dMRI experiments with a range of b-values where the signal decay is still mono-exponential (typically *b* ≤ 1.2 *ms/µm*^2^) as *S*(*b*) = *S*(0) exp(−*b* · *ADC*). Here, a maximum b-value of 1 *ms/µm*^2^ was used and ADC was estimated using nonlinear-least squares within a mask derived at each time point. Organoid volume was estimated from the binary mask at each time point, by multiplying the number of voxels included in the mask with the voxel volume.

##### Diffusion kurtosis imaging

Young and mature organoids were scanned with trapezoidally-shaped diffusion gradients with fixed Δ and *δ* (Table 1, kurtosis experiment), but with variable gradient strengths to achieve b-values up to 3 *ms/µm*^2^. All other parameters, including TE, were kept fixed. ADC and kurtosis *K* were estimated using nonlinear-least squares as *S*(*b*) = *S*(0) exp(−*b* · *ADC* + *b*^2^ · *ADC*^2^ · *K/*6).

##### ADC diffusion-time dependence

Organoids were scanned with trapezoidally-shaped diffusion gradients of varying diffusion time (Δ − *δ/*3), achieved by varying Δ at a constant *δ* (Table 1, diffusion-time dependence experiment). dMRI data were acquired for a range of b-values up to 1 *ms/µm*^2^ to estimate ADC. All other parameters were kept fixed.

##### Diffusion tensor imaging

Organoids were scanned with trapezoidally-shaped diffusion gradients with fixed Δ and *δ* (Table 1, diffusion tensor imaging experiment), b-values up to 1 *ms/µm*^2^, and 9 gradient orientations uniformly distributed over the sphere. All other parameters were kept fixed. The diffusion tensor **D** was estimated using ordinary least squares in FSL as *S*(*b*, **g**) = *S*(0) exp(*b***g**^*T*^ **Dg**). The eigenvectors and eigenvalues of the diffusion tensor were derived, and mean diffusivity 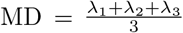 and fractional anisotropy 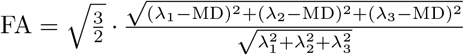 computed. Streamline tractography was performed on the first eigenvector – indicating the main direction of diffusion. The minimum FA for tracking was set to 0.2, the maximum angle to 30°, step size to half the voxel size.

##### Frequency-dependent diffusion tensor-relaxation distributions

Organoids were scanned with a protocol comprising diffusion encoding waveforms beyond standard trapezoidal shapes (i.e. tensor-valued diffusion encoding) (Figure 1 (b)) to resolve distributions of diffusion tensors instead of one macroscopic diffusion tensor per voxel obtained in the protocol of the diffusion tensor imaging section. Specifically, these acquisitions enable the disentangling of heterogeneity in (an)isotropy and time-dependence within a voxel, using 0 and first-order modulation waveforms [64, 90]. Additionally, variable *TE* and *TR* were acquired to assess heterogeneity in relaxation times, by introducing variable echo time delays (*τ*_*E*+_) and repetition time delays (*τ*_*R*+_) (Figure 1 (b)) to achieve a given *TE* and *TR* without changing the diffusion-encoding timings and enhance relaxation contrast. The acquisition protocol designed for this study was composed of 307 images with *TR* ranging from 0.92 to 3.42 *s, TE* from 16 to 46 *ms*, b-values from 0.03 to 4.07 *ms/µm*^2^, with waveform-duration from 3 to 9 *ms*, the encoding anisotropy *b*_Δ_ of −0.5, 0, and 1, and centroid frequency *ω*_*cent*_ from 50 to 300 *Hz* for *b*-weighted images. The *ω*_*cent*_ were estimated as the centroid of the gradient moment power spectrum [95]. The imaging parameters were a FOV (4.3 *mm*)^3^, matrix (72)^3^, spatial resolution (60 *µm*)^3^, EPI readout bandwidth 132,086 *Hz*, 1 segment, leading to a total experiment time of 15 *h* 18 *min*.

Non-parametric distributions of tensor-valued diffusion spectra **D**(*ω*), where **D** are axially symmetric positive semi-definite diffusion tensors parameterised by frequency-dependent axial and radial diffusivities *D*_∥_(*ω*) and *D*_⊥_(*ω*), and first eigenvector direction (Θ, Φ))) and longitudinal *R*_1_ and transversal *R*_2_ relaxation rates, were estimated using Monte Carlo inversion of the multidimensional dataset according to the following equation [64, 67]:

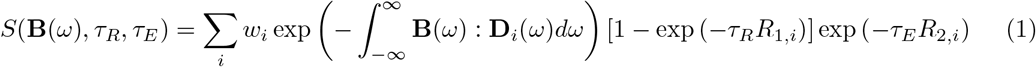

The frequency-dependent diffusivities were further parameterised as

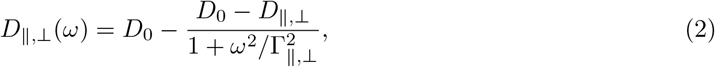

where *D* are the corresponding zero-frequency values, *D*_0_ the intrinsic diffusivity at high-frequency, and Γ the frequencies at the centers of the transitions. Thus, each discrete component *i* in the **D**(*ω*) − *R*_1_ − *R*_2_ distribution can be described by the parameters [*w, D*_∥_, *D*_⊥_, *D*_0_, Γ_∥_, Γ_⊥_, Θ, Φ, *R*_1_, *R*_2_].

The Monte Carlo inversion was performed with Matlab using the *md-mri* Toolbox with the limits: 5 ·10^−12^ *m*^2^*s*^−1^ *< D*_0,∥,⊥_ *<* 5 ·10^−9^ *m*^2^*s*^−1^, 10^−1^ *s*^−1^ *<*Γ_∥,⊥_ *<* 10^5^ *s*^−1^, 0.1 *s*^−1^ *< R*_1_ *<* 4 *s*^−1^, and 4 *s*^−1^ *< R*_2_ *<* 200 *s*^−1^, 20 steps of proliferation, 20 steps of mutation/extinction, 200 input components per step of proliferation and mutation/extinction, 10 output components. The bootstrapping was performed by 100 repetitions using random sampling with replacement to compute the parameter maps. Specifically, means *E*, variances *V* and covariances *C* over dimensions were computed within a voxel. The frequency dependent isotropic diffusivity *D*_iso_ and diffusion anisotropy *D*_Δ_ maps were created by evaluating *D*_∥,⊥_(*ω*) at *ω* = 51 *Hz* and computing *D*_iso_ = (*D*_∥_ + 2*D*_⊥_)*/*3 and *D*_Δ_ = (*D*_∥_ − *D*_⊥_)*/*3*D*_iso_. Frequency dependence maps were created by

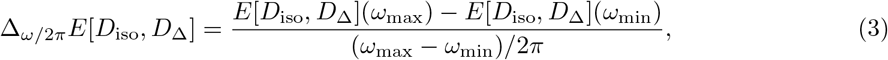

with *ω*_max_ = 214 *Hz* and *ω*_min_ = 51 *Hz*.

The parameter correlation allows for separating the distributions into subsets (bins) based on the value of a chosen parameter, and obtaining the distribution of values of all other parameters in each subset[96]. A three bin’s model defined based on the *D*_*iso*_ and *D*_Δ_^2^ was suggested for brain-like structures to discriminate white matter (WM), gray matter (GM) and cerebrospinal fluid (CSF) [64, 68]. According to this model, bin 1 highlights the low diffusivity and high anisotropy components and corresponds to the WM compartment. Bin 2 shows low diffusivity and low anisotropy components corresponding to the GM-like compartment. Bin 3 represents high diffusivity compartments corresponding to CSF. For the cortical brain organoids, only bin 1 and 2 were analyzed. The metrics *D*_*iso*_ and *D*_Δ_^2^ were calculated for the low frequency *ω* = 51 Hz. Specifically, the bin-boundaries were defined according to: bin1: *D*_*iso*_ *<*1·10^−9^ *m*^2^*s*^−1^ and 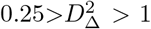; bin2: *D*_*iso*_ *<*1·10^−9^ *m*^2^*s*^−1^ and 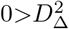 *>* 0.25; and bin3: *D*_*iso*_ ≥ 1· 10^−9^ *m*^2^*s*^−1^. To avoid extrapolation, the frequency-dependent parameters (*D*_*iso*_ and 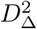) are quantified at frequencies restricted to the 10 and 90 percentiles of the *ω* values sampled, corresponding to 51 and 214 *Hz*.

### 3.3 Microscopy

Following MRI measurements, organoids underwent a fixation preparation procedure. The culture medium was removed from the MR glass tubes and replaced with PBS, filling the tubes to the top. The organoids were then carefully transferred to Eppendorf tubes pre-filled with PBS. This was accomplished by inverting the MR glass tube over the Eppendorf tube, allowing the organoid to gently sink to the bottom of the Eppendorf. Subsequently, the PBS in the Eppendorf tube was replaced with 4 % PFA to initiate fixation (Fisher Scientific, 50-980-487) in 1× PBS (Life Technologies, 14190169) for either 2 hours at room temperature (RT), or overnight at 4 ^*o*^*C*. Subsequently, organoids were washed 3 times with 1× PBS and stained according to the protocol below.

The iDISCO clearing procedure was performed as described [97, 98]. In short, perfused, isolated, and post-fixed brains were gradually dehydrated in a series of 20 % MeOH (VWR, Cat*#*1.06009.2500), 40 % MeOH, 60 % MeOH, 80 % MeOH, and 100 % MeOH (2×), for 1 *h* at RT each, followed by overnight incubation (ON) at RT in 66 % DCM (Sigma-Aldrich, Cat*#*5895810250) and 33 % MeOH. The next day, samples were incubated in 100 % MeOH (2×) for 1 *h* at RT, followed by incubation at 4^*o*^*C* for more than 1 *h* (to cool the sample for bleaching). Next, samples were bleached ON in 5 % hydrogen peroxide (VWR, Cat*#*1072100250) in MeOH at 4 ^*o*^*C*. The following day, samples were gradually rehydrated using a series of 80 % MeOH, 60 % MeOH, 40 % MeOH, 20 % MeOH for 1 *h* each at RT. Samples were then washed with 1× PBS for 1 *h* at RT. Subsequently, samples were incubated twice in Ptx.2 (1× PBS and 0.2 % Triton X-100) for 1 *h* at RT and transferred into permeabilization solution (80 % Ptx.2, 20 % DMSO, 2.3 % glycine) for 24 *h* at 37 ^*o*^*C* on a horizontal shaker (70 *rpm*). This shaker was also used for blocking and antibody incubations with the same settings. Samples were incubated in blocking solution for 4 days, followed by incubation in primary antibody solution (PwtH (1× PBS, 0.2 % Triton X-100, 0.1 % Heparin (Sigma, Cat*#*H3393, 10 mg/mL)) stock solution, 5 % DMSO and 3 % normal donkey serum (NDS) (Jackson Immunoresearch, Cat*#*017-000-121)) for 3 days at 37 ^*o*^*C* while shaking (each day a fresh antibody solution was added). Then, samples were washed 5 times for 1 *h* at RT in a 15 *mL* Falcon tube containing PtwH. Samples were incubated in secondary antibody solution (PwtH and 3% NDS (Jackson Immunoresearch, Cat*#*017-000-121)) for 1 day at 37 ^*o*^*C* on a horizontal shaker (70 rpm) in a 5 *mL* Eppendorf. Primary antibodies used, were mouse anti ZO-1 1:250 (Bdbiosciences, Cat*#*610966) in combination with secondary donkey anti-mouse IgG Alexa Fluor 488 1:750 (Thermo Scientific, Cat#A21202), SMI312 1:500 (BioLegend, mouse, Cat*#*837904,) in combination with secondary donkey anti-mouse IgG Alexa Fluor 1:750 (Thermo Scientific, Cat#ab175738). TO-PRO3 (Thermo Scientific, Cat*#*T3605) was added in the mix of secondary antibodies to stain the nucleus of cells in the far red spectrum. Initially, the concentration for TO-PRO3 was 1:5000, and it was later increased to 1:1000. Before the tissue clearing, samples were embedded in 1 % agarose (Sigma-Aldrich Cat*#*11388983001) in TAE. Then, samples were dehydrated using a series of 20 % MeOH, 40 % MeOH, 60 % MeOH, 80 % MeOH and 2 times 100 % MeOH, each step for 1 *h* at RT. Samples then were placed in 66 % DCM and 33 % MeOH for 3 *h* at RT, followed by 2 washes in 100 % DCM for 15 *min* at RT each. After removing lipids with Dichloromethane (DCM), clearing was finalized by incubation in Dibenzylether (DBE) (Sigma-Aldrich, Cat*#*108014). For clearing, tissue samples were placed in 100% DBE overnight at RT. Samples were stored in 100 % DBE at RT until imaging. All washing, dehydration and clearing steps were performed in dark Falcon tubes to protect against light and on a rotator (14 rpm). The organoids were imaged with an UltraMicroscope Blaze (Miltenyi Biotec) lightsheet microscope equipped Neo sCMOS camera (2048×2048 pixels. Pixel size: 6.5 x 6.5 *µm*^2^) and Imspector software (version 7.7. (Miltenyi Biotec)). Samples were images with12x NA 0.53 MI PLAN objective and 0.6x zoom via the magnification changer, the objective contained a dipping cap for organic solvents. The sample was imaged with single sided illumination, a sheet NA of 0.135 (results in a 4 *µm* thick sheet) and a step-size of 2 *µm* using the horizontal focusing lightsheet scanning method with optimum amount of steps and using the fixed blend algorithm. The effective magnification was 7.2 (magnification changer *objective = 0.6*12). The following filter combinations were used: Ex: 500/20 + Em: 595/40, Ex:560/40 + EM: 620/60, Ex: 630/30 + EM:680/30, Ex:785/25 + EM:805LP in combination with a white light superK Fianium laser FIU-15 (NKT Photonics). Nuclear density of cortical organoids was estimated from nuclei counts and organoid volumes measured in Imaris (Version 10.2.0). Nuclei counting was performed with an estimated nucleus diameter of 3 µm, background subtraction, and detected spots filtered by the quality metric with a threshold above 158. Organoid volumes were obtained by manually drawing a mask around each organoid and using the Imaris surface function (no background subtraction, surface grain size of 10 µm) with manual thresholding. Volumes were measured from both 3D LS and diffusion-weighted MRI images, and the volumes differed, most likely due to staining procedures. Reported nuclear densities in this paper are based on volumes measured from a 3D dMRI experiment.

## 4 Acknowledgements

CMWT and TN are supported by a Vidi grant (21299) from the Dutch Research Council (NWO), and this research was funded in part by the Wellcome Trust [215944/Z/19/Z]. MY is supported by the Fondation pour la Recherche sur le Cancer (ARC). For the purpose of Open Access, the author has applied a CC BY public copyright licence to any Author Accepted Manuscript version arising from this submission. We thank Itamar Ronen, Andrew Webb, Willeke van Roon-Mom, Ronald Buijsen, Aldrik Velders and Alia A for valuable discussions and Andrei Gurinov and Johan van der Zwan for technical assistance. Technical support from Bruker is acknowledged.

This research was supported by uNMR-NL Grid, an NWO-funded National Roadmap Large-Scale Facility of the Netherlands (projects 184.032.207 and 184.035.002).

